# Context-dependent epigenome rewiring during neuronal differentiation

**DOI:** 10.1101/2024.10.18.618996

**Authors:** Vera Manelli, Jeisimhan Diwakar, Sude Beşkardeş, Dácil Alonso-Gil, Ignasi Forné, Faye Chong, Axel Imhof, Boyan Bonev

## Abstract

Transcription factors (TFs) are pivotal in orchestrating lineage decisions and their binding is often accompanied by chromatin remodeling. However, a comprehensive understanding of how cellular and epigenetic contexts influence TF binding and the subsequent activation of lineage-specific programs remains lacking. Here, we dissected the molecular mechanisms underlying the binding of one such TF – Neurog2 and its functional consequences in two cell-types: mouse embryonic stem cells and neural progenitor cells. Our findings reveal that cell-type-specific chromatin accessibility and motif syntax are key determinants for TF binding. While Neurog2 binding primarily leads to chromatin opening, DNA demethylation and increased chromatin interactions, we also uncovered strong indirect, cell-type-specific effects, which ultimately result in vastly different epigenetic landscape. Furthermore, we identify shared and cell-type-specific Neurog2 interactors, including the SWI/SNF and NuRD complexes. Our study shed light on how cellular environment can modulate TF function to establish lineage identity during development.

## Introduction

Development is characterized by progressive differentiation and specification into a multitude of distinct cell types, which requires finely tuned spatiotemporal gene regulation. During this process, various transcription factors (TFs) directly engage with regulatory elements in the genome, allowing them to activate or repress gene expression networks and establish cell type-specific transcriptional programs^1^. Notably, despite the presence of thousands of transcription factor binding motifs (TFBMs) across the genome, only a small percentage becomes occupied, even in the case of transcription factors that can interact with inaccessible chromatin^2^. Epigenetic regulation converging on chromatin accessibility or DNA methylation^3–5^ has been shown to contribute to TF binding selectivity but the precise molecular mechanisms are still not fully understood. Furthermore, most studies that examined TF binding have focused only on a single cell type, and it is unclear to what extend the genomic and cellular contexts contribute to this process.

In addition to being important for TF selectivity, epigenetic remodeling has also been observed downstream of TF binding. Such rewiring can occur across multiple modalities such as DNA methylation^6,7^, chromatin accessibility^7,8^, 3D genome rewiring^9–11,7^, and frequently involve interactions with chromatin remodelers. This can be leveraged to induce cell differentiation^12^, trans-differentiation^13,14^, and even de-differentiation^6^. Indeed, various elements have been predicted to contribute to their activity, such as more or less permissive chromatin states, the availability of other interaction partners, the presence of different isoforms, or the presence of post-translational modifications (PTM)^15^. However, a major unresolved question is how TFs can accomplish such widespread remodeling and, importantly, to what an extent such changes depend on the genomic and cellular contexts.

Direct neuronal differentiation or reprogramming is ideally suited to address these questions as it represents a fate change elicited by a single TF. One of the most potent TFs in this paradigm is Neurog2, which has been used to derive induced neurons from mouse and human pluripotent stem cells^16^, as well as to reprogram differentiated cell types such as astrocytes^14,17^ to neurons. Neurog2 is a TF of the basic helix-loop-helix (bHLH) family controlling the activation of various lineage programs responsible for the development of the central nervous system^18–20^. These include the regulation of progenitor cell proliferation and patterning, alongside orchestrating neuronal differentiation, subtype specification, and migration.

Neurog2 binding has been associated with increased chromatin accessibility during fibroblast reprogramming^21^, embryoid bodies^22^, developing mouse cortex and astrocytes^7,14^. Additionally, gain-of-function experiments *in vivo* have shown that the presence of Neurog2 leads to DNA demethylation and the formation of regulatory loops^7^. Based on this data, we have previously proposed that Neurog2 acts as a “molecular bridge” by coordinating changes across multiple epigenetic layers to allow robust and irreversible commitment to the neuronal lineage^7,14^.

Neurog2 is typically expressed transiently during development and previous work suggests that the developmental time window and cell type in which Neurog2 acts can influence the identity of the neurons that are subsequently generated^17,23^. Indeed, Neurog2 overexpression has been leveraged in generating various neuronal populations such as glutamatergic, motor, sensory, dopaminergic, and serotoninergic neurons *in vitro*^16,24–29^, but the cellular mechanisms driving this specificity are unclear. Thus, the study of the controlled expression of Neurog2 in different cellular contexts represents a general framework to dissect master TF-induced epigenomic changes across distinct cell types.

To achieve this goal, here we employed a doxycycline-inducible mouse embryonic stem (ES) cell line to precisely and controllably overexpress Neurog2 in both ES and differentiated neural progenitor cells (NPC). We found both shared and cell type-specific chromatin binding patterns and identified motif syntax as essential in determining chromatin remodeling at Neurog2-occupied regions. Neurog2 binding generally leads to increased chromatin accessibility, DNA demethylation and stronger chromatin contacts in both cell types. However, Neurog2 also indirectly results in genome-wide increased DNA methylation, decreased accessibility at sites occupied by REST and influences global 3D genome organization. Finally, we showed that the direct effects are likely mediated via interactions with chromatin remodellers and the NuRD complexes, while indirect effects are likely due to ectopic binding and subsequent misregulation due to the cellular and epigenetic context. Thus, our study represents a framework for comprehensively investigating the molecular mechanism and the functional importance of cell-type-specificity in TF mediated lineage decisions.

## Results

### Neurog2 induces both shared and cell-type-specific transcriptional response

To investigate the influence of cellular context for Neurog2-mediated epigenome rewiring, we first generated a mouse ES line containing a Flag-Neurog2 doxycycline-inducible construct (Tet-ON system) from a safe harbour locus, that allows tightly controlled expression in ES and NPCs (Fig. 1a-b). We then induced Neurog2 expression with 24hrs dox treatment in ES cells as well as differentiated NPCs. Surprisingly, we observed higher Neurog2 protein levels in ES, while mRNA was stronger in NPC (Fig. 1b and Extended Data Fig. 1c-f). We next asked if Neurog2 overexpression resulted in changes of cellular identity. We observed a slight reduction of pluripotency-associated markers in ES and an overall increase in neuronal markers such as ßIII-tubulin (Tubb3). However, NPC markers such as Pax6 were largely unchanged (Fig. 1c and Extended Data Fig. 1b). Overall, Neurog2 induction was highly reproducible and resulted in 3707 differentially expressed genes (DEGs) in ES and 3072 DEGs in NPC (Fig. 1d and Extended Data Fig. 1g).

**Figure 1.**
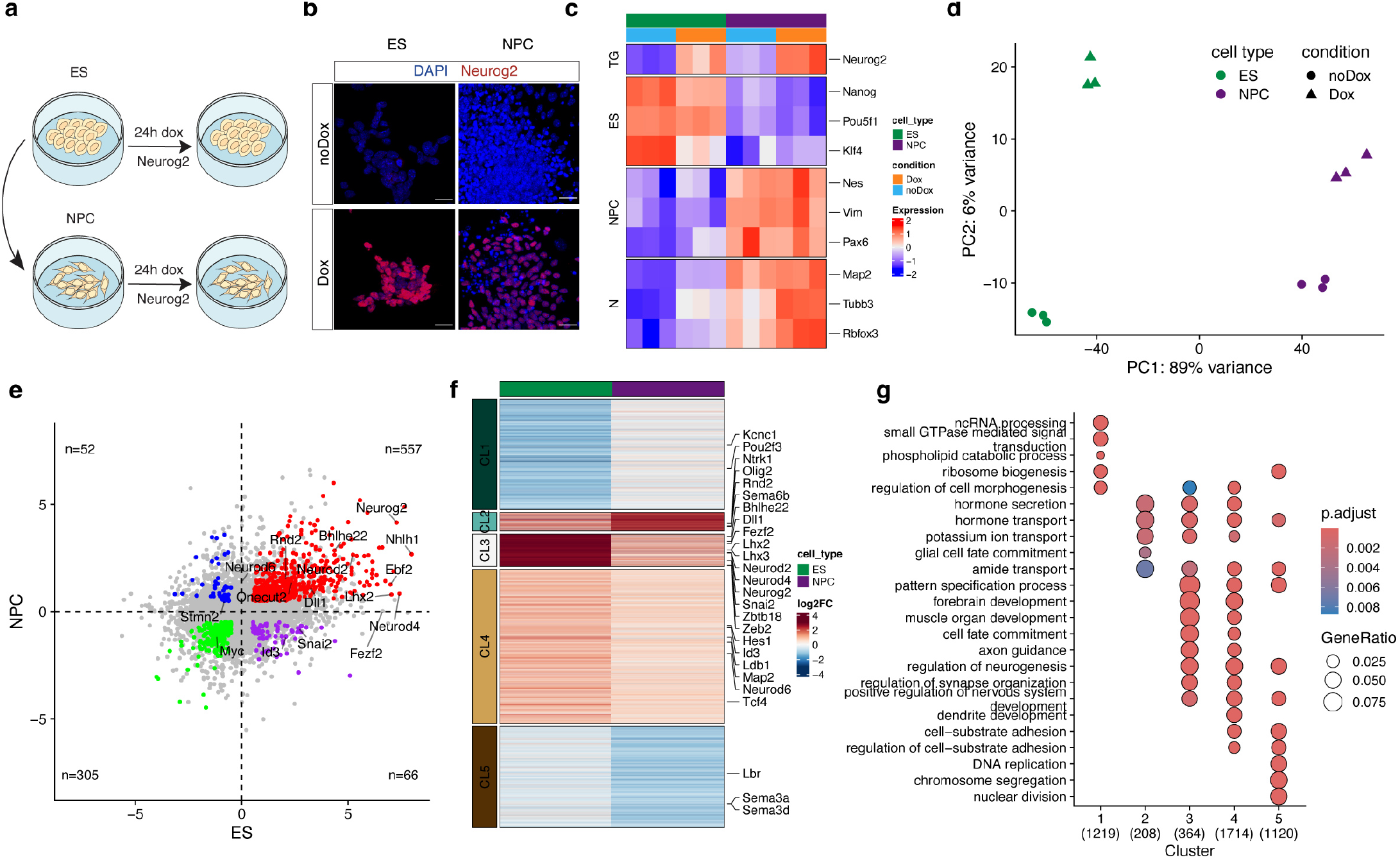
Neurog2 drives cell-type-specific and conserved transcriptional responses. (**a**) Schematics of the experimental strategy. The inducible ES cell line was either supplemented with doxycycline for 24 hours or differentiated to NPC and subsequently supplemented with doxycycline for 24 hours. (**b**) Representative immunostaining of Neurog2 in ES and NPC with and without doxycycline. (**c**) Heatmap displaying normalized expression of ES, NPC or neuronal (N) markers upon differentiation and/or Neurog2 induction (n=3) (d) PCA analysis of the transcriptomic data. (e) Direct 4-way comparison of transcriptional changes across cell types and conditions. Colored dots represent significantly upregulated (red), downregulated (green) or anti-correlated genes (blue and purple). (f) K-means clustering (k=5) of differentially expressed genes in at least one of the two cell types. (g) Dot plot depicting the gene ratios of the enriched gene ontology terms across the identified RNA-seq clusters in (f). The color of the circles represents the Benjamini–Hochberg adjusted P value.

Next, we asked if the transcriptional programs activated by Neurog2 are shared between the two cell types. We observed that the majority of DEGs were correlated between both cell types with a smaller proportion being specific only to one cell type (Fig. 1e). Among the commonly upregulated genes (n=557), there were previously reported direct targets such as Dll1, Rnd2, Lhx2, Neurod2, Neurod1 and Neurod4^7,17,20,22,30^, together with Tcf4 and other members of the bHLH TF family such as Bhlhe22 and Nhlh1^31^, involved in the development of mouse nervous system (Fig. 1e). NPC-specific upregulated genes include Stmn2, encoding for a microtubule-stabilizing protein involved in the neurite outgrowth^32^, and Neurod6, which is generally expressed in a subset of differentiated pyramidal neurons located in the deep layers of the cortex^33^. Conversely, we identified Id3, which forms heterodimers with bHLH TFs and prevents their binding to DNA^34^ to be specifically upregulated in ES, but not in NPC (Fig. 1e).

To further identify common and cell type-specific transcriptional programs, we clustered all DEGs identified in at least one of the two cell types (Fig. 1f). Genes upregulated in NPC but not in ES were mostly enriched for small GTPase-mediated signaling pathway (cluster 1, Fig. 1f-g), while ES specific genes were involved in DNA replication, chromosome segregation, and nuclear division (cluster 5, Fig. 1f-g). DEGs with stronger upregulation in NPCs were associated with ion transport and glia cell fate (cluster 2, Fig. 1f-g), while most of the known Neurog2 target genes involved in neurogenesis were in cluster 3 (n=364), typically upregulated stronger in ES compared to NPCs. However, even within this cluster the absolute levels of expression varied (Extended Data Fig. 1h). To account for these differences in abundance, we also performed a direct comparison between DEGs in ES and NPC, which identified transcriptional repressors such as Id3 and Snai2, genes that encode for other members of the bHLH family like Tcf12, and Sox9, implicated in Sox2 inhibition in development^35^ as ES-specific. We also observed that ES-specific genes were associated with alternative differentiation lineages such as renal or mesenchymal cells (Extended Data Fig. 1j).

Taken together, these findings suggest that Neurog2 orchestrates changes in transcriptional networks to drive neural fate commitment. Although most target genes are consistently up- or down-regulated across both cell types, a small but significant subset is cell-type-specific. These finding point to cellular and genomic context as an important aspect of Neurog2-mediated changes in gene expression.

### Cell-type-specific Neurog2 binding occurs at already accessible regions

To understand how Neurog2 mediates these transcriptional changes, we performed ChIP-Seq in both cell types and identified 1946 ES-specific, 2537 NPC-specific and 15199 shared peaks (Fig. 2a). Neurog2 binding was stronger in ES, even at shared peaks (Extended Data Fig. 2a), consistent with the higher protein levels (Extended Data Fig. 1e-f).

**Figure 2.**
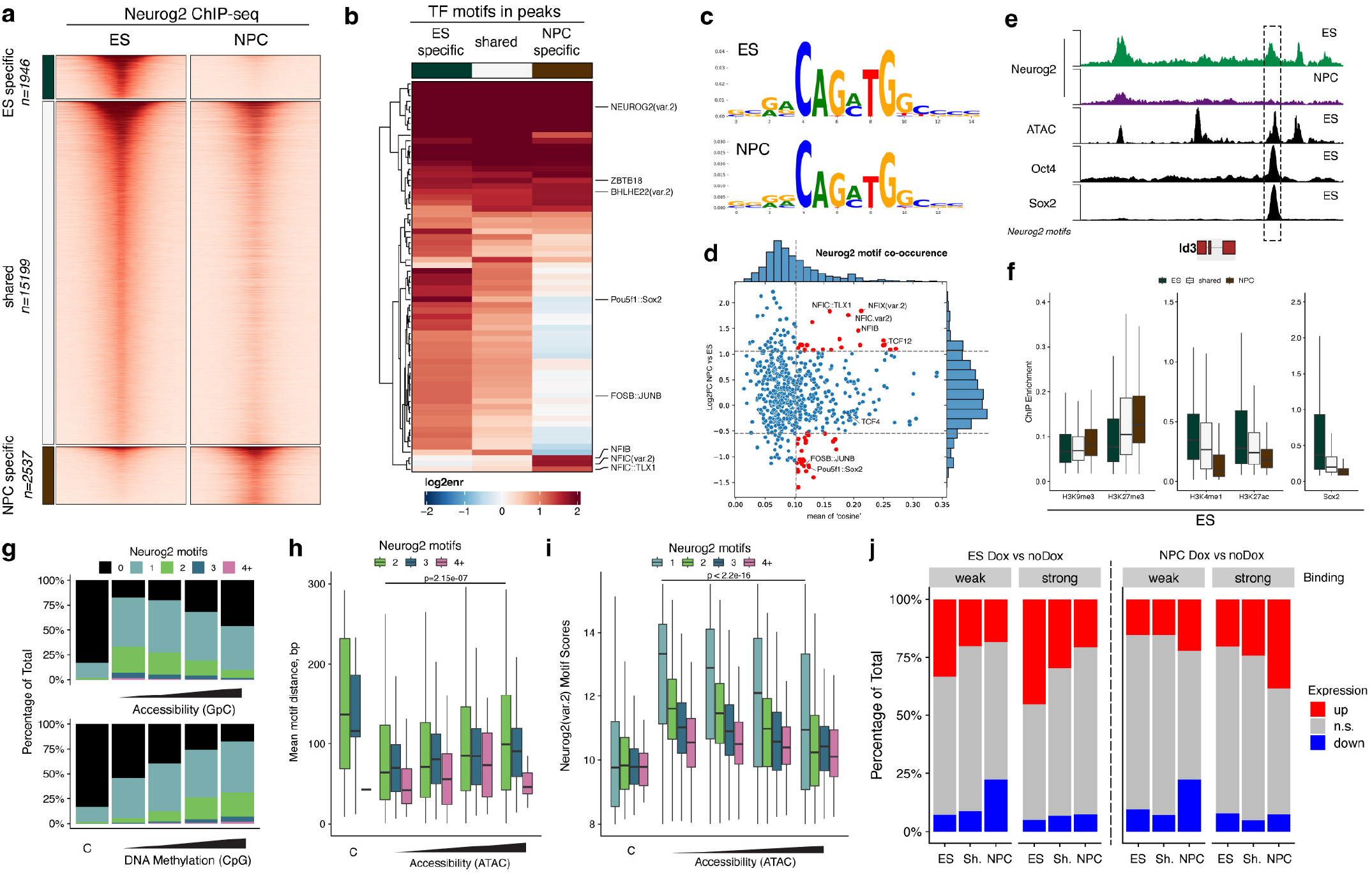
Cell-type-specific and independent Neurog2 binding. (a) Heatmap displaying Neurog2 ChIP-Seq enrichment at shared or differentially bound peaks in ES and NPCs. (b) Heatmaps depicting TF motif enrichment analysis in the peak groups displayed in (a). (c) Contribution weight matrix (CWM) for the most significant motif in both ES and NPC. (d) Motif co-occurrence comparing mean cosine similarity and fold change across motif pairs containing Neurog2. (e) Example genomic tracks depicting Neurog2, Oct4 and Sox2 ChIP-seq tracks as well as ATAC at the Id3 locus. ES-specific putative enhancer co-bound by Neurog2, Oct4 and Sox2 is highlighted. (f) Boxplots depicting the ChIP-seq enrichment in reads per million (RPM) of the various ES histone marks^11^ at the indicated peak categories (ES = ES-specific, NPC = NPC-specific, shared = present in both ES and NPCs) (g) Percentage of Neurog2 peaks with a given number of motifs at either control regions or Neurog2 shared ChIP-seq peaks, stratified by pre-existing GpC-based accessibility or CpG DNA Methylation in ES^39^ (h) Boxplot showing the distribution of the motif distances within a peak stratified by accessibility (i) Boxplot showing the distribution of Neurog2 motif scores within a peak stratified by accessibility (j) Percentage of differentially regulated genes, whose promoters are bound by Neurog2 in ES and NPC. Genes are additionally split in two quantiles based on Neurog2 binding (weak or strong).

We then compared binding profiles to previous studies and found that Neurog2 binding in ES was similar to embryoid bodies 12h after induction^22^, while the ones in NPCs were more similar to the 48h timepoint and to the *in vivo* Neurog2 (Extended Data Fig. 2b)^36^.

Next, we asked which TF-binding motifs were enriched in each peak group. Both shared and cell-type specific peaks were enriched in the previously characterized Neurog2 motif (Fig. 2b). To understand if subtle motif variations could explain the differential binding, we performed both kmer enrichment and deep learning analysis, based on BPnet and TFMoDisco^37^. Both approaches identified the canonical E-box signature CAG-ATG, suggesting that Neurog2 was bound to the same motif in both ES and NPC (Fig. 2c and Extended Data Fig. 2c). We then asked if the presence of other TF motifs could explain the cell-type-specific binding. We observed that ES-specific peaks were enriched in Oct4-Sox2 motifs, while NPC-specific peaks displayed an increased number of motifs of TFs from the Nfi family, involved in cortical differentiation^38^ (Fig. 2b, d and Extended Data Fig. 2e). The presence of Sox2 and Nfi at the prospective cell-type specific Neurog2 peaks was also confirmed by ChIP-seq (Extended Data Fig. 2d). These results are exemplified at the *Id3* locus, which is upregulated specifically in ES (Fig. 1e). Here we observed Neurog2 binding consistent with Id3 expression at several distal regulatory regions which were accessible and occupied by Oct4/Sox2 but had no Neurog2 motif (Fig. 2e).

We then asked if Neurog2 cell type-specificity could be preferentially associated with different chromatin features prior to binding. ES-specific peaks were also enriched for H3K27ac and H3K4me1 in ES, while NPC-specific peaks were associated with the repressive mark H3K37me3 (Fig. 2f and Extended Data Fig. 2f). These results were also consistent with nucleosome occupancy, showing that ES-specific peaks were located primarily at nucleosome-free regions, while shared and especially NPC-specific peaks have higher nucleosome occupancy in ES (Extended Data Fig. 2g).

Since motif distance and cooperativity have previously been shown to influence pioneer TFs^37^, we focused on their contribution to Neurog2 binding, stratified by pre-existing chromatin accessibility and DNA methylation levels in ES cells^39^. Importantly, we observed that the binding at lowly accessible and highly methylated regions correlated with the presence of multiple motifs, suggesting a synergistic effect (Fig. 2g and Extended Data Fig. 2h). Shorter motif distances and motif scores were also predictors of binding at closed chromatin (Fig. 2h-i).

Finally, to investigate the relationship between Neurog2 binding and transcriptional activity, we compared the percentage of differentially regulated genes that overlap with shared or cell-type-specific peaks. Neurog2 binding in both ES and NPCs preferentially led to gene activation (Fig. 2j), This effect was more prominent at cell-type specific peaks and was correlated with the binding strength (Fig. 2j). This was exemplified at the *Id2* locus, where Neurog2 binding at its promoter is cell type-specific and results in ES-specific transcriptional upregulation (Extended Data Fig. 2i).

Overall, these results suggest that multiple features such as pre-existing chromatin accessibility, presence of other TFs, motif number and/or distance modulate the affinity of pioneer TFs such as Neurog2 to chromatin, leading to cell-type specific binding and transcriptional regulation.

### Direct and indirect effects of Neurog2 binding on chromatin accessibility

Having established the importance of genomic and epigenetic features for Neurog2 binding specificity, we next asked what the effect on the epigenome is downstream of Neurog2. To accomplish this, we employed 3DRAM-Seq, a multi-omics method that measures genome-wide chromatin accessibility alongside DNA methylation and 3D genome organization^39^. To avoid any secondary effects from changes in the cell cycle, we selected G_0_/G_1_ cells in both conditions. Furthermore, to avoid any potential indirect effects due to inefficient differentiation or cell fate switch for NPC, we additionally purified only Pax6^high^ population which we refer to just NPC for simplicity (Extended Data Fig. 3a).

We first focused on changes in chromatin accessibility and uncovered unexpected cell-type specific differences. We observed that in NPC, Neurog2 binding at both distal and promoter regions was mostly associated with increased accessibility, consistent with its role as a pioneer TF (Fig. 3a, top). This effect was stronger at NPC-specific peaks and resulted in increased accessibility at the Neurog2 motifs (Fig. 3b-c, top). Conversely, we observed that in ES many Neurog2-bound regions were also associated with decreased accessibility, which was especially prominent at promoters and ES-specific peaks (Fig. 3a-b, bottom) but not at Neurog2 motifs (Fig. 3c, bottom), suggesting that loss of accessibility could be a potential indirect effect.

**Figure 3.**
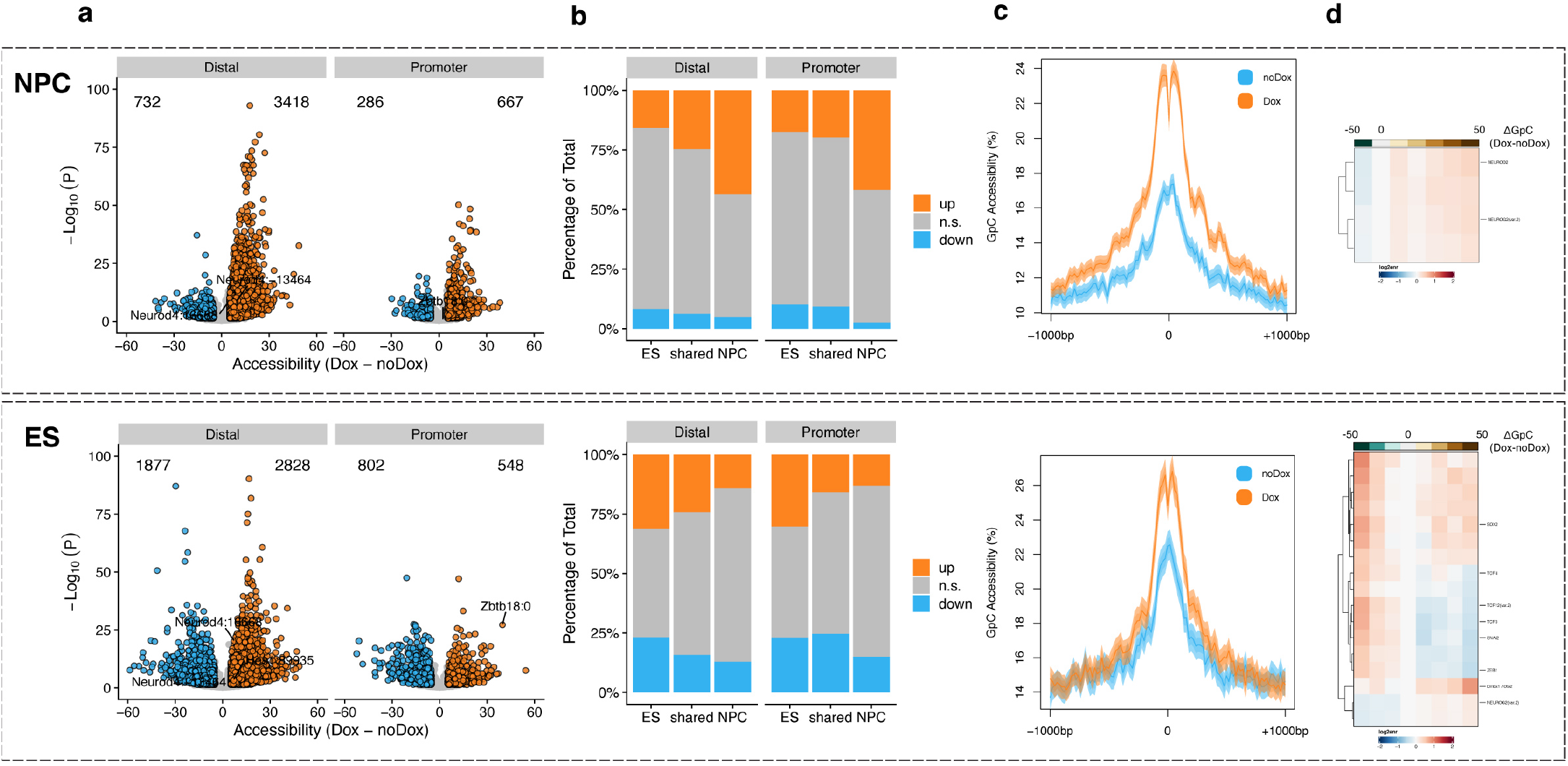
Neurog2 binding leads to increased chromatin accessibility. (a) Volcano plots displaying differentially accessible Neurog2 peaks at distal and promoter regions upon Neurog2 overexpression. (b) Percentage of differentially accessible Neurog2 peaks across different peak categories. (c) Neurog2 motif footprinting (in top 5000 distal peaks). (d) Heatmap depicting TFBM enrichment for Neurog2 peaks in ES (upper panel) and NPCs (lower panel) stratified by the changes in accessibility

Using motif enrichment analysis, we found that regions which become more accessible, were enriched predominantly in Neurog2 motifs in both ES and NPC (Fig. 3d and Extended Data Fig. 3d). In addition, homeobox TFs such as Otx2 which have been linked to enhancer activation^40^ were also enriched only in ES, which was correlated with its cell-type specific upregulation (Extended Data Fig. 3f). Conversely, we identified E proteins and Ebox-binding transcriptional repressors such as Snai2 as enriched in Neurog2 peaks which lose accessibility specifically in ES. These findings suggest that Ebox repressors compete with Neurog2 for binding at the same regions and higher levels in ES (Extended Data Fig. 3g) can lead to loss of accessibility.

However, E proteins and Ebox repressors alone were not sufficient to explain the overall loss of chromatin accessibility observed in ES (Extended Data Fig. 3b-c, bottom). Therefore, we examined TF motifs in regions associated with loss of accessibility genome-wide and identified REST as highly enriched in this category (Extended Data Fig. 3d). This could result from lower REST occupancy at these sites, increased repression or both. To discriminate between these hypotheses, we performed motif footprinting and observed lower accessibility and less nucleosome phasing in ES and to a lesser extent also in NPC (Extended Data Fig. 3e). This was consistent with the significant decrease in REST expression (Extended Data Fig. 3h) and upregulation of REST target genes (Extended Data Fig. 3i), indicative of overall release from REST-mediated repression.

Taken together, these findings suggest that Neurog2 remodels chromatin accessibility in two separate ways. Its binding directly leads to increased chromatin accessibility both in ES and NPC. However, there is also considerable cell-type specificity upon Neurog2 expression (primarily in ES) - increased compaction mediated locally by Ebox proteins such as Snai2 and genome-wide due to reduced REST occupancy.

### Neurog2 is associated with increased global methylation in ES

Next, we asked if Neurog2 also leads to changes in DNA methylation, as previously reported^7^. Indeed, we observed that many distal regions bound by Neurog2 were demethylated upon Dox induction in both ES and NPC, while the effect on promoters was less obvious (Fig. 4a). This effect was strongest in NPC-specific peaks and resulted in decreased methylation at Neurog2 motifs (Fig. 4b-c, top). To our surprise, we observed that some Neurog2 peaks also become methylated in ES, which was especially prominent at distal regions (Fig. 4a-b, bottom). However, similar to the effects on chromatin accessibility, Neurog2 motif was primarily demethylated (Fig. 4c, bottom), pointing to a potential indirect effect. Indeed, we observed increased DNA methylation levels genome-wide in ES, including gene bodies and distal regions (Extended Data Fig. 4a-b).

**Figure 4.**
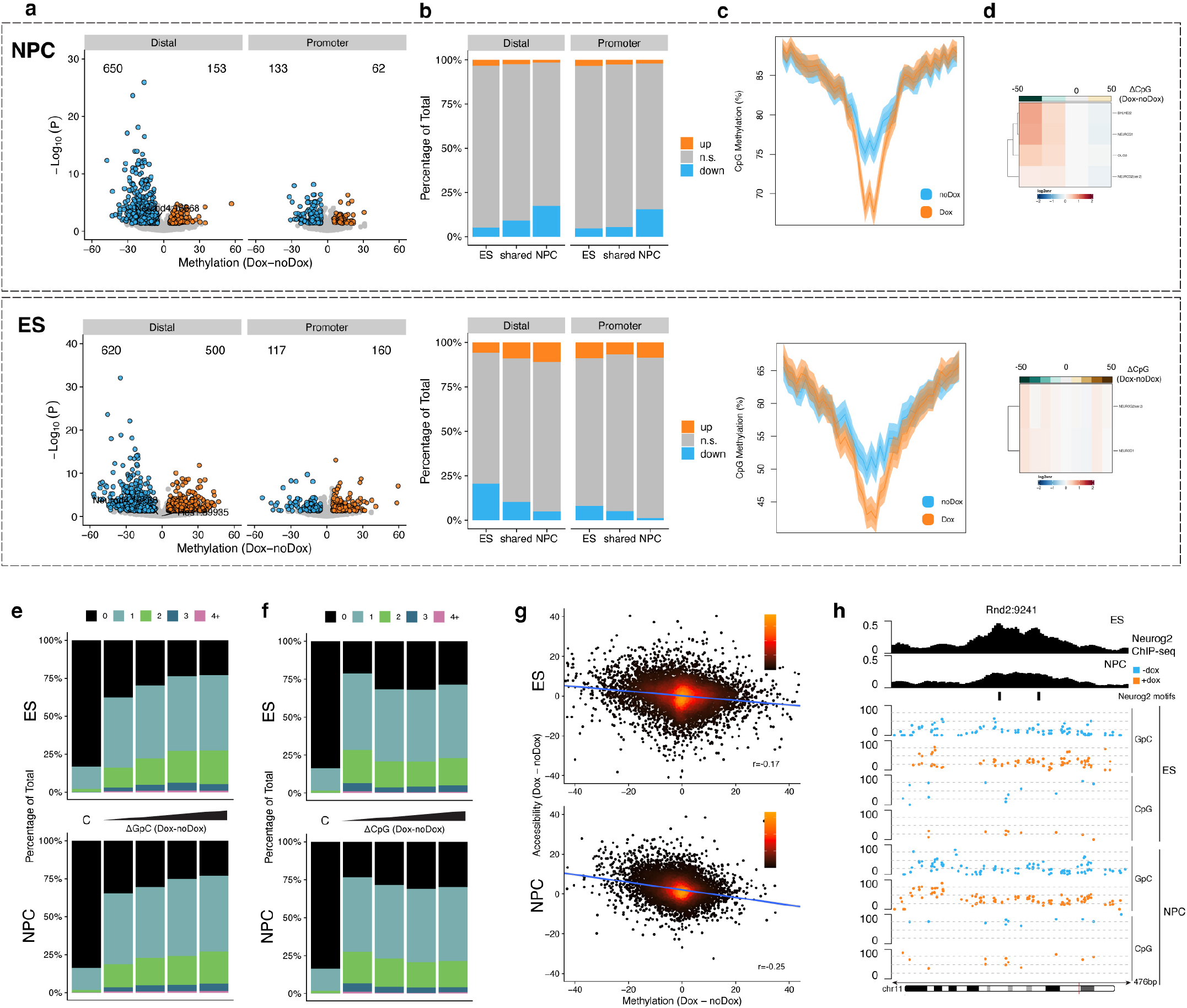
DNA methylation dynamics and correlation to chromatin accessibility at Neurog2 peaks. (a) Volcano plots displaying differentially methylated Neurog2 peaks at distal and promoter regions upon Neurog2 overexpression. (b) Percentage of differentially methylated Neurog2 peaks across different peak categories. (c) DNA methylation profile at Neurog2 motifs within top 5000 distal peaks. (d) Heatmap depicting TFBM enrichment for Neurog2 peaks in ES (upper panel) and NPCs (lower panel) stratified by the changes in DNA methylation. (e-f) Percentage of Neurog2 peaks with a given number of motifs at either control regions or Neurog2 shared ChIP-seq peaks, stratified by the change in chromatin accessibility (e) or DNA methylation (f) in ES or NPC (g) Density plots displaying changes of CpG methylation and GpC accessibility at Neurog2 peaks in ES or NPC. Spearman’s correlation is at the lower right corner (h) Example genomic tracks depicting Neurog2 ChIP-seq, Neurog2 motifs, CpG DNA methylation and GpC chromatin accessibility tracks at the *Rnd2* locus.

To further understand these changes, we performed motif enrichment analysis and observed only bHLH motifs and especially Neurog2 were enriched in the regions that become demethylated (Fig. 4d and Extended Data Fig. 4c), while methylation at REST sites was mostly unchanged (Extended Data Fig. 4c). Surprisingly, such an effect seemed to be reversed on the global scale in ES, where most differentially methylated regions (DMRs) were associated with increased methylation levels instead. This widespread increase in DNA methylation in ES was correlated with downregulation of Tet1 levels and transcriptional upregulation of Dnmt3a (Extended Data Fig. 4e-f)^41^, where we observed ES-specific binding of Neurog2 at a putative intragenic regulatory site (Extended Data Fig. 4g).

Finally, we asked if changes in chromatin accessibility and/or DNA methylation are also correlated with Neurog2 binding. We observed that sites that became more open or demethylated upon Neurog2 binding tended to have a higher proportion of co-occurring Neurog2 motifs (Fig. 4e-f). Interestingly, the anticorrelation between changes in accessibility and DNA methylation levels was overall weak, especially in ES (Fig. 4g), suggesting that these two modalities could be partially uncoupled.

### Neurog2 leads to direct and indirect 3D genome rewiring

Having examined chromatin accessibility and DNA methylation, we next asked how Neurog2 affects 3D genome organization. We sequenced 4 billion reads in total and observed high correlation across replicates (Extended Data Fig. 5a) and good QC indicative of high-quality libraries (Table 1).

Focusing first on Neurog2 binding sites, we observed an increase in contact frequency both in ES and NPC (Fig. 5a-b). To address if this is due to recruitment of cohesin at the Neurog2 peaks, we performed Rad21 Cut&Run and observed increased enrichment at those regions (Fig. 5c), which was more pronounced in the ES compared to NPC.

**Figure 5.**
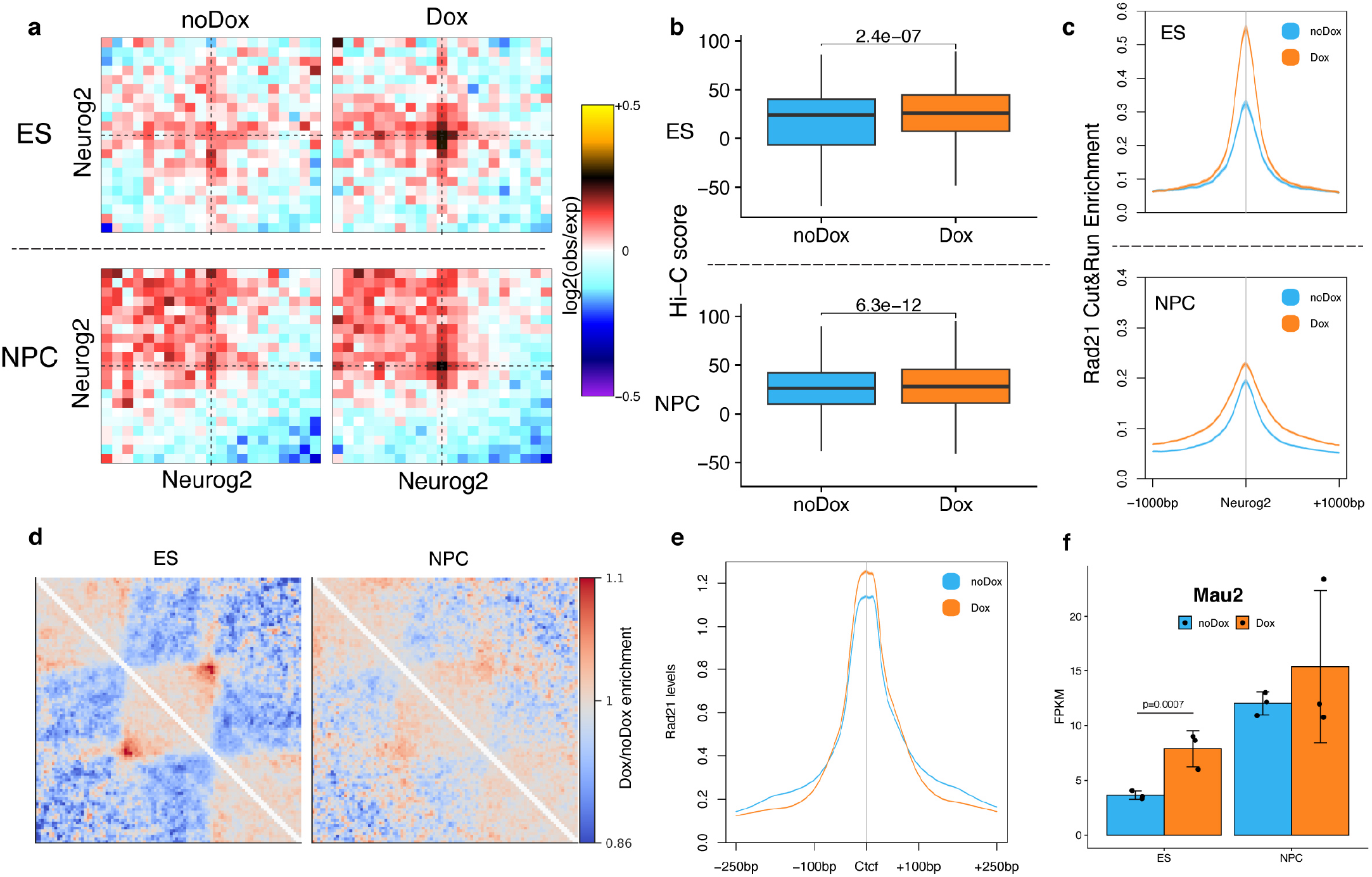
Neurog2 leads to both direct and indirect 3D genome rewiring. (a) Aggregated Hi-C plots between intra-TAD pairs of the top 5000 Ngn2 ChIP-seq peaks. (b) Boxplot displaying the Hi-C score of Neurog2 binding sites in ES and NPC (top and bottom). Statistical significance is calculated using paired Wilcoxon test. (c) Rad21 enrichment at Neurog2 ChIP peaks in ES and NPC (d) Fold change in chromatin contacts at TADs in ES or NPC (e) Rad21 enrichment at Ctcf motifs within Ctcf ChIP-seq peaks in ES (f) Expression of *Mau2* in either ES or NPC (dots represent individual biological replicates, error bars represent SD). Statistical significance is derived from Deseq2 Wald test.

Next, we asked whether Neurog2 is also associated with genome-wide changes as we had previously observed at the linear epigenome scale. We identified a shift towards short-range contacts upon dox induction only in ES but not in NPC (Extended Data Fig. 5b). This was not due to changes in compartmentalization, as compartment strength appears to be mostly unchanged between the two conditions (Extended Data Fig. 5c-d). In contrast, we observed that chromatin insulation and interactions between TAD boundaries were increased specifically in ES but not in NPC (Fig. 5d and Extended Data Fig. 5e). This effect was also present more generally at Ctcf sites, where both insulation and looping were increased in ES (Extended Data Fig. 5f-g).

Next, we hypothesized that altered cohesin levels on chromatin, binding or processivity could underlie these results. To address this, first we measured Rad21 levels on chromatin using immunoFACS but didn’t observe any major differences not related to the cell cycle (Extended Data Fig. 5h). Furthermore, most of Rad21 binding sites were present in both conditions (Extended Data Fig. 5i) and the dox-specific peaks were mostly associated with Neurog2 binding (Extended Data Fig. 5j). However, when we examined the distribution of Rad21 around Ctcf sites, we observed that it became sharper – i.e. higher maximum enrichment while lower dispersion (Fig. 5e). These results suggest that cohesin is more efficiently “blocked” from extruding loops at Ctcf sites. Consistent with this, we also observed that the levels of Mau2 (also known as Scc4), which together with Nipbl is required for cohesin loop extrusion^42^, were increased upon Neurog2 expression specifically in ES but not in NPC (Fig. 5f).

Taken together these results suggest that Neurog2 affects 3D genome organization in two separate ways. First, its binding leads to increased looping in both ES and NPC, likely due to increased cohesin levels at Neurog2 sites. However, Neurog2 expression also has profound effects on global 3D genome organization specifically in ES: increased compaction, stronger insulation and looping genome-wide, which is likely indirect and could arise from more efficient blocking of loop extruding cohesin by Ctcf due to increased Mau2 levels.

### Neurog2 is associated with known bHLH TFs and chromatin remodelers

To understand the differences in the cell-type specific epigenetic remodelling, we decided to examine potential co-factors and interactors of Neurog2. Therefore, we profiled proteins associated with Neurog2 (either directly or indirectly via chromatin) using ChIP-mass spectrometry (ChIP-MS) in both ES and NPC (Fig. 6a).

**Figure 6.**
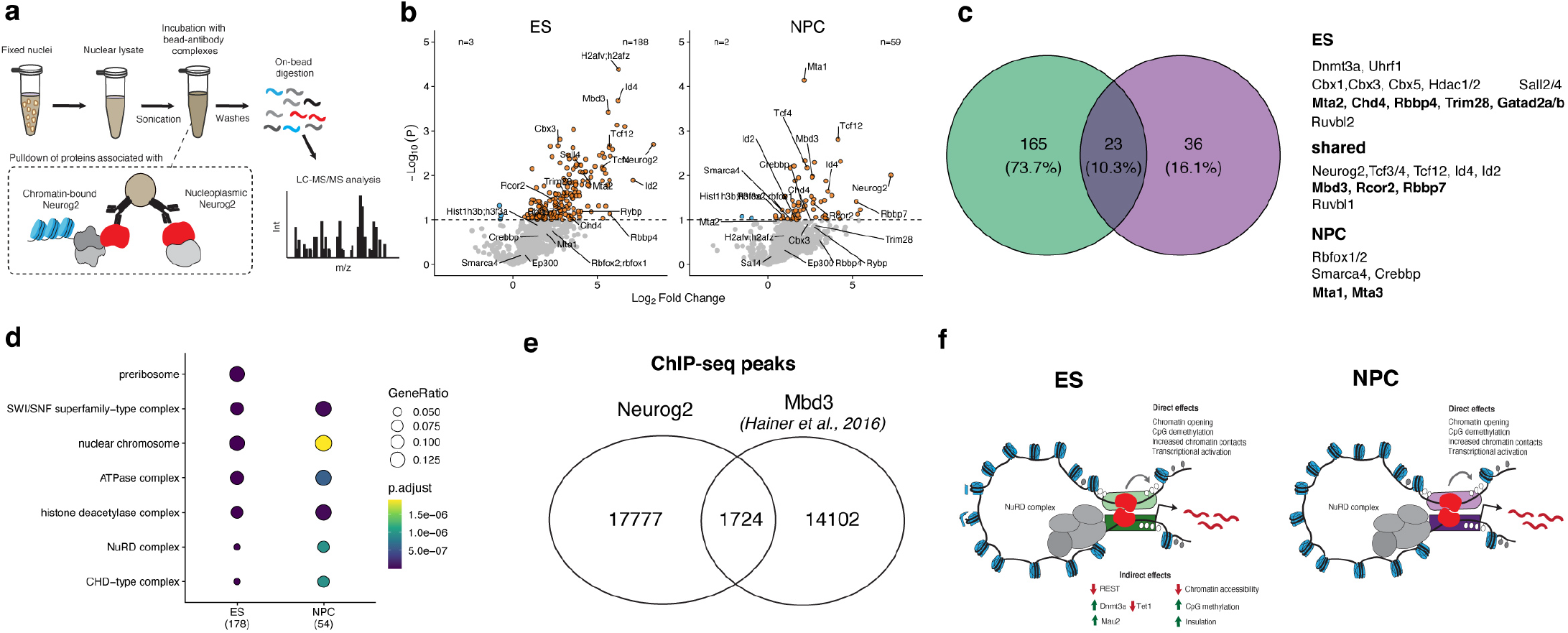
Neurog2 interacts with chromatin remodelers and the NuRD complex. (a) Schematics of ChIP-MS for Neurog2-associated proteins (b) Volcano plot displaying representative proteins associated with Neurog2 identified through ChIP-MS analysis (n=3 replicates). (c) Venn diagram displaying the shared and cell type-specific significantly enriched Neurog2 -associated proteins. (d) Dot plot displaying GO terms enrichment in the hits from (b). (e) Venn diagram displaying the percentage of overlap between Neurog2 peaks and published Mbd3 ChIP-seq peaks in ES (Hainer et al., Elife 2016) (f) schematic representation of the main findings of the study.

We identified 224 proteins significantly enriched in the Neurog2 ChIP-MS data, with 10.3% of the hits present in both ES and NPC (Fig. 6b-c). Neurog2 itself, as well as other E-box factors known to heterodimerize with Neurog2, such as Tcf3/4/12 and Id proteins^43^, were identified in both cell types, confirming the efficiency and the specificity of the approach.

In addition to E-box proteins, we also identified several chromatin-remodeling complexes (Fig. 6c-d). Among them, the most prominent was the NuRD complex, with multiple subunits such as Mbd3, Rbbp7 and Rcor2^44^ among the hits shared in both ES and NPC. Other components of the NuRD complex were found to interact in a cell-type-specific manner: Mta2, Rbbp4, Trim28, Gatad2a/b (p66α/β) and Chd4^44–46^ were present in ES, while Mta1/3 were found to interact only in NPC. Other repressive complexes which have been linked to NuRD such as Hdac1/2 and Sall2/4 were also specifically observed in ES but not in NPC. In addition to the NuRD complex, we also detected interactions with Ruvbl1 (Rvb1) – a core subunit of the INO80 complex, which is associated with nucleosome remodeling^47^. Furthermore, we observed proteins associated with transcriptional activation such as the major SWI/SNF subunit Brg1 (Smarca4) and Crebbp (Cbp) interacting with Neurog2 specifically in NPC but not in ES. Importantly, neuronal Brg1 phosphorylation has been reported to affect enhancer-promoter loops and influence its interaction with the NuRD complex^48^ and may contribute to its cell-type specific interaction with Neurog2 only in NPC.

Finally, we asked if the interaction with the NuRD complex could be due to Neurog2 binding to regions already occupied by NuRD. To address this, we compared Neurog2 and Mbd3 bound regions but found that most sites do not overlap (Fig. 6e), suggesting that pre-existing NuRD binding cannot explain the interactions with Neurog2.

Overall, these results suggest that Neurog2 coordinates epigenetic rewiring by interacting with both E-box proteins and chromatin remodelers. Some of those interactions are shared (such as with key subunits of the NuRD and the Ino80 complex), while others are cell-type specific (such as the SWI/SNF and Cbp) pointing to the importance of cellular context for transcription factor-mediated cell fate changes.

## Discussion

Here, we used two different cellular systems to uncover the shared and context-specific determinants of transcription factor binding and its downstream effects on chromatin accessibility, DNA methylation, 3D genome organization and ultimately gene expression. Furthermore, we linked these widespread remodeling at the epigenetic landscape to shared and cell-type specific interacting partners which include chromatin remodeling complexes and histone variants linked to enhancer regulation. Thus, we comprehensively characterize the molecular mechanism of how transcription factors can both read and reshape the cellular and epigenetic context to change the fate of the cell.

Previous approaches to study the molecular mechanism of pioneer TFs have typically focused on cases where the TF is endogenously expressed^49^ or used heterogenous systems undergoing differentiation^22^. Although such systems have led to valuable insights, they have mostly relied on correlation between binding and chromatin state or have only focused on chromatin accessibility without examining additional epigenetic layers. Importantly, almost all studies have focused on a single cell type, thus making it difficult to uncover the contribution of cellular and epigenetic context to TF binding and downstream function.

We chose to focus on two cell types which can be used to address how TFs function either in “optimal” environment, such as Neurog2 in NPC (where it normally is expressed during development), or in cells which are highly plastic, but it is not normally expressed (such as ES). We observed differences in the amount of Neurog2 protein levels, which could be due to posttranslational regulation. This is consistent with previous reports of complex interplays of phosphorylation and ubiquitination affecting Neurog2 stability^50–53^. Furthermore, Neurog2 phosphorylation at different residues has been suggested to influence Neurog2 proneural activity both in development^43,54^ and in reprogramming^14^. Therefore, it is possible that the differential PTMs could also contribute to both cell-type-specific binding and downstream chromatin remodeling, eventually leading to differential gene regulation.

We observed that the majority of the genes regulated by Neurog2 become upregulated in both cell types, consistent with its previously described role as a transcriptional activator^30^. However, we also identified many genes which were differentially regulated in only one condition, including more than 100 which were induced in one but repressed in another cell type, such as Id3 and Snai2. Since these effects were present already by 24h, it is tempting to speculate that they may further contribute to the divergence of the lineage programs activated by Neurog2 in different cell types – a phenomenon observed in reprogramming^17^.

Differential cellular and genomic context has been speculated to also influence the binding of pioneer TFs, but the potential mechanisms are unclear. Here, we found that pre-existing differential accessibility affects the binding of TFs such as Neurog2, even if they can bind to initially closed chromatin regions. These results are consistent with a recent preprint showing that the number of Neurog2 motifs and their distance are correlated with binding at closed regions and changes in accessibility in ES cells^55^. Therefore, even though most binding sites are shared across cell types, around 20% are cell-type specific and are often associated with binding of other TFs such as Oct4/Sox2 in ES, and the Nfi family of TFs in NPC. Thus, the resulting ectopic binding at open regions can lead to misregulation of genes such as Id3 and further divergent lineage programs due to indirect effects. This is important conclusion for the field of reprogramming as limiting such ectopic effects, for example by targeted closing of specific binding sites via CRISPRi, can potentially improve conversion efficiency.

Consistent with previous reports, we observed that Neurog2 binding is generally associated with gene activation and increased chromatin accessibility^7,14,22^. This is often accompanied by DNA demethylation at Neurog2-bound distal regions, however, there are many regions where the dynamics of the two epigenetic modalities (chromatin accessibility and DNA methylation) are uncoupled. Finally, we also observed increased chromatin interactions between two Neurog2 bound sites, which is likely due to the increased recruitment of cohesin upon Neurog2 binding. Interestingly, another bHLH protein, MyoD has recently been shown to directly promote the formation of chromatin loops^56^. Therefore, we propose that formation of chromatin loops between lineage defining TFs could be a general mechanism to ensure co-regulation of target genes and ensure the robustness of the response. It remains to be seen if such properties are limited to just bHLH proteins or could be generalized to other key TFs.

In addition to these direct effects, we also uncovered an unexpected influence of the cellular context on the epigenetic rewiring downstream of Neurog2 (Fig. 6f). In ES, there is a global decrease of accessibility, especially at sites bound by Rest and the pluripotency complex Oct4/Sox2. We believe that this is likely indirect and it is due to the downregulation of REST itself as well as the decrease in expression of the pluripotency factors. This indirect change in chromatin accessibility is also accompanied by a genome-wide increase in DNA methylation levels. In addition to accessibility and DNA methylation, we also observed genome-wide increase in short-range interactions and chromatin insulation, but not compartment strength in ES. This is accompanied by stronger enrichment of Rad21 at Ctcf sites, which we propose is at least partially due to the upregulation of one of the proteins implicated in cohesin sliding/processivity – Scc4 (Mau2). Our findings are consistent with recent studies showing that depletion of Scc4 has the opposite effect – weaker/shorter loops and reduced chromatin insulation^57^.

Finally, using ChIP-MS we identified prominent interactions between Neurog2 and chromatin remodeling complexes. In particular, the association with subunits of the SWI/SNF and NuRD complexes may clarify some of the observed changes in nucleosome positioning and chromatin remodeling. Although NuRD is traditionally associated in repression^58^, recent evidence indicates that NuRD subunits are found at nearly all active enhancers and promoters in ES cells, with some of its subunits specifically linked to the active transcriptional machinery^59^. This suggests that the association of the NuRD complex with Neurog2 may extend beyond the passive co-occupation of chromatin and play an active role in the fine-tuning of the targeted developmental genes. Alternatively, Neurog2 might recruit the NuRD complex to repress alternative cell fates similar to what has been reported recently for other pioneer TFs such as FoxA1^60^. We hypothesize that such repressive interactions are more likely to occur in ES, which are much more plastic and primed to adopt alternative lineages. Along those lines, we observed cell-type-specific interactions with the repressive proteins Sall2/4 specifically in ES, which could indicate shared molecular mechanisms with other lineage TFs such as FoxA^60^.

Taken together, our multi-modal dissection of the effects of TF action on the various epigenetic layers has uncovered that lineage defining TFs such as Neurog2 can have both shared as well as cellular and epigenetic context-dependent properties. This phenomenon is observed both upstream, in that pre-established accessibility contributes to cell-type specific binding, but also modulates the molecular cascades downstream of the TF. Here we uncovered that extensive, yet most likely secondary, epigenome rewiring events are more pronounced in ES, the cell type that is more distant from the cellular context in which Neurog2 generally operates. Collectively, these findings lay the groundwork for delineating the causal relationship among the epigenetic changes at the core of local and global TF-driven events, shedding light on novel aspects of reprogramming and differentiation in different cellular environments.

## Author Contributions and Notes

V.M., J.D and B.B. conceptualized the study. V.M. performed the RNA-seq, ChIP-seq and ChIP-MS and contributed to the data analysis. J.D. performed the 3DRAM-seq experiments. S.B performed the Cut&Run experiments. D.A-G performed the chromatin Rad21 experiments. F.C contributed to the data analysis. I.F and A.I assisted with the MS data analysis. B.B assisted with experiments, analyzed data and supervised the project. V.M and B.B wrote the manuscript with input from all authors. The authors declare no competing interests.

## Acknowledgments

We thank Prof Magdalena Götz, Prof Gunnar Schotta and all members of the Bonev lab for useful discussions. Sequencing was performed at the Helmholtz Zentrum München by staff at the NGS-Core Facility and cell sorting was done at the Flow Cytometry Core Facility at the Biomedical Center, Ludwig-Maximilians-Universität. V.M. was supported by a Boehringer Ingelheim Fonds (BIF) PhD Fellowship. B.B. was supported by Helmholtz Center Munich, DFG priority programme SPP2202 (BO 5516/1-1), ERA-NET Neuron (MOSAIC) and European Research Council Consolidator grant (EpiCortex, 101044469).

## Materials and Methods

### Cell culture

MEFs (Gibco, Cat.N:A34181) were plated on 0.1% gelatin-coated plates (Merck, Cat.N ES-006-B) as per the manufacturer’s instructions and supplemented with DMEM (Thermofisher, Cat.N: #21969-035) containing 10% FBS (Thermofischer, Cat.N:16141079), 50U pen-strep (Gibco, Cat.N: 16141079), 0.1 mM of non-essential amino acids(Gibco, Cat.N: 11140035), and 0.1 mM of 2-mercaptoethanol (Gibco, Cat.N: 31350010). The medium was changed every other day.

Flag-Neurog2 ES cell lines were maintained as described in Bonev et al., 2017^11^, with some changes. In brief, cells were cultured on MEFs in DMEM (ThermoFisher, Cat.N:21969-035), supplemented with 15% FBS (Thermofischer, Cat.N:16141079) 1,000 U/mL of LIF (Millipore, Cat.N:ESG1107), 0.1 mM of non-essential amino acids (Gibco, Cat.N:11140035), 50U of pen-strep (Gibco, Cat.N:15140-122) and 0.1 mM of 2-mercaptoethanol (Gibco, Cat.N:31350010). Media was changed daily and 3×10^5 cells were passaged every other day using TryplE (Life technologies, Cat.N: 12604013) with a 30-minute selective sedimentation step where the detached cells were kept in an uncoated 10cm dish at 37C to separate ES from MEFs. For Neurog2-overexpression, cells were detached and replated without MEFs. Doxycycline was supplemented on the following day and cells were harvested after 24 hours, unless otherwise specified.

### Flag-Neurog2 inducible cell line generation

A2Lox.cre mES cell lines^61^ were supplemented with 500 ng/mL doxycycline for 24 hours. After detachment and counting, 1×10^6 cells were subjected to electroporation (Amaxa nucleofector 96-well shuttle, solution VHPH-1001, waveform program 96-CG-104) with 1 μg of p2Lox-Flag-Neurog2 plasmid, followed by plating on neomycin-resistant MEFs (Gibco, Cat.N: A34963). Selection with 300 μg/mL G418 (Gibco, Cat. N: 11811023) was initiated 24 hours after plating and was continued for 10 days. Colonies were manually picked on day 10, then replated and expanded on neomycin-resistant MEFs.

### Neural differentiation

Cells were differentiated into neuronal progenitors as described^11^ with minor changes. In brief, cells were plated at low density (1×10^5 cells per plate) on gelatin-coated 10cm dishes (without MEFs) in ES media and after 12h cultured in DDM media (DMEM/F12 + GlutaMAX (ThermoFisher, Cat.N: 31331-028), supplemented with 1x N2 (ThermoFisher, Cat.N: 17502-048), 1 mM of sodium pyruvate (ThermoFisher, Cat.N: 11360-070), 500 ug/ml BSA Fraction V (ThermoFisher, Cat.N:15260-037), 50U of pen-strep (Gibco, Cat.N:15140-122) and 0.1 mM of 2-mercaptoethanol (Gibco, Cat.N:31350010). for a total of 12 days. Cyclopamine (Merck, Cat. N.: 239803) was added at 1uM concentration every two days from the second day to day 10 of differentiation. Media was changed every two days. Doxycycline was supplemented on day 11 and cells were harvested upon dissociation with Trypsin after 24 hours, unless otherwise specified.

### Immunocytochemistry

ES and NPCs were grown on 0.1% gelatin-coated coverslips in 12-well plates. After 2 washes in 1x DPBS, they were fixed in 4% methanol-free Formaldehyde (Pierce, Cat.N: 28908) in PBS for 20 minutes at room temperature and washed two times in cold 1x DPBS. Successively, the coverslips were incubated with a permeabilization solution (0.3% Triton X-100 in 1xDPBS) for 20 minutes at RT and then with the blocking solution (BSA fraction V 3% (ThermoFisher, Cat.N:15260-037), 0,3% TritonX-100, 5% horse serum (Sigma, Cat.N: H0146-10ML) in 1xDPBS). Subsequently, the coverslips were immersed in the antibody dilution buffer (0,1% TritonX-100, 1% horse serum in 1xDPBS) containing the primary antibody of interest **(See Antibody section)** and were left incubating overnight at 4C. The day after, they were washed 3x with washing buffer - 0.1% Triton X-100 in PBS for 10 mins each at RT, followed by the incubation with the secondary antibody of choice (see Antibody section) in the antibody dilution buffer.

The second wash contained DAPI diluted 1:1500. After the last wash, the coverslips were kept in 1x DPBS and mounted individually by using Prolong Diamond (Thermofisher, Cat.N: P36961). The imaging happened within the following 15 days. The acquisition of the images was carried out with an LSM710 laser-scanning confocal and Zen2 software (Carl Zeiss).

### Antibody list for Immunocytochemistry

#### Primary antibodies

| Name | Species | Cat.N: | Dilution |
| --- | --- | --- | --- |
| Pax6 | Rabbit | 901301 | 1:100 |
| beta tubulin | Mouse | 560394 | 1:400 |
| Nanog | Mouse | 560259 | 1:200 |
| Neurog2 | Mouse | MAB3314 | 1: 100<br>(stock: 0.5 mg/mL in PBS) |

#### Secondary antibodies

|  | Species | Cat # | Dilution |
| --- | --- | --- | --- |
| Anti-rabbit (488) | Donkey | A32790 | 1:1000 |
| Anti-rabbit (555) | Donkey | A32794 | 1:1000 |
| Anti-rabbit (647) | Donkey | A31573 | 1:1000 |
| Anti-mouse (488) | Donkey | A32766 | 1:1000 |
| Anti-mouse (555) | Donkey | A32773 | 1:1000 |
| Anti-mouse (647) | Donkey | A31571 | 1:1000 |

### Bulk RNA-Seq

RNA was extracted from ES and NPCs as follows. Upon harvesting, approximately 5×10^5 cells were pelleted and resuspended in Trizol (Thermofischer, Cat.N: 15596026). After 5 minutes at room temperature samples were vortexed for 20 s, 0.2x volumes of chloroform was added, tubes were mixed by inverting and samples were centrifuged at 13,000 rpm at 4C for 15min. The aqueous phase was then processed using the Zymo kit (Zymo research, Cat.N: R1013) with DNase treatment, according to the manufacturer’s instructions. 100ng of RNA per each condition was used as a starting material for cDNA synthesis and library preparation with the Low Input RNA Library Prep Kit for Illumina® (New England Biolabs, Cat.N: E6420). The quality of the RNA-seq libraries was assessed using the Agilent 2100 Bioanalyser system (Agilent).

### Protein extraction for western blot

Cells were resuspended in cell lysis buffer (150 mM NaCl, 1% Igepal, 0.5% sodium deoxycholate, 0.1% SDS, 50 mM Tris, pH 8.0, 50mM NaF, 1x protease inhibitor Roche) for a total of 30 minutes on a rotating wheel at 4C and centrifuged at maximum speed at 4C. The supernatant was conserved at -20C and used within the following days.

### Western Blot

The protein samples (within a range of 6-10ug) were first incubated with Pierce™ Lane Marker Reducing Sample Buffer (Cat.N: #3900) and denatured at 95°C for 5 minutes and left on ice for 10 minutes.The proteins were separated on a Precast Mini-PROTEAN TGX Gels Gradient 4-20% (Novex, Cat.N: 4561083) in running buffer (200mM glycine, 25 mM Tris, 3.5 mM SDS) using a voltage of 100V. Subsequently, the proteins were transferred to nitrocellulose membranes (Abcam, Cat.N: ab133413) using transfer buffer (200mM glycine, 25 mM Tris, 1.5 mM SDS) at 100V for 1 hour at 4°C. After the transfer, the membranes were blocked in TBST (10 mM Tris, 100 mM NaCl, pH 7.4, 0.1% Tween-20) with 5% (W/V) non-fat milk for 1-2 hours at room temperature, supplemented with primary antibodies and subsequently incubated overnight at 4°C. The membranes wee then washed three times for 10 minutes each in TBST and incubated for 1 hour at room temperature with secondary antibodies (See antibody list) Following another three washes for 10 minutes each at room temperature with TBST, the membranes were processed using Forte (Thermofischer, Cat.N: WBLUF0100) before being imaged using the Biorad Chemidoc MP imaging system.

List of antibodies used for western blot

Primary antibodies

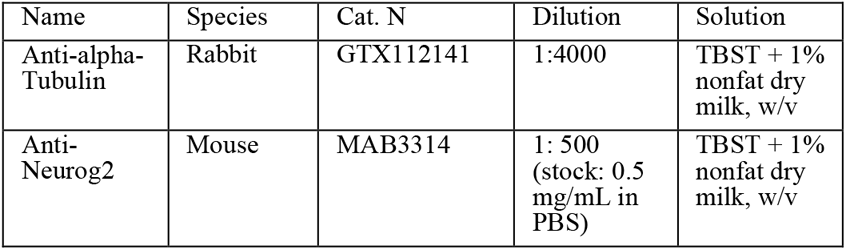

Secondary antibodies:

| Name | Species | Cat. N | Dilution | Solution |
| --- | --- | --- | --- | --- |
| Ant-Rabbit-HRP | Goat | 111-035-144 | 1:4500<br>(previously diluted 1:1 glycerol) | TBST + 1% nonfat dry milk, w/v |
| Anti-Mouse-HRP | Goat | 111-035-003 | 1:4500<br>(previously diluted 1:1 glycerol) | TBST + 1% nonfat dry milk, w/v |

### Quantification of Western blot bands

The quantification Neurog2 levels was achieved via ImageJ (FIJI). The corrected integrated of density of Neurog2 was normalized to the corrected integrated density of alpha tubulin in the same lane to account for variations in sample loading and efficiency of transfer.

### Chromatin Immunoprecipitation

The ChIP-seq protocol was adapted from Bonev et al., 2017 ^11^. Briefly, 10×10^6 cells were fixed using 1% methanol-free Formaldehyde (Pierce, Cat.N: 28908) at room temperature for 10 minutes at a concentration of 1×10^6 cells/ml and subsequently quenched with glycine at a final concentration of 0.2M, followed by incubation for 5 minutes at room temperature. Both fixation and quenching were carried out with gentle rotation on a shaker (40 rpm). Fixed cells were snap-frozen in liquid nitrogen and kept at -80C.

All steps were performed at 4C and all the bead washes were preceded by the tubes being on a magnet for 1 minute, unless otherwise specified. The cells underwent lysis for 10 minutes in 500ul cold lysis buffer (10mM Tris pH 8.0, 0.2% NP-40, 10 mM NaCl) supplemented with 1x cOmplete EDTA-free protease inhibitor (Roche, Cat.N:05056489001), followed by a spin at 500xg for 5minutes, one wash in cold 1xDPBS and another spin under the same conditions. Nuclei were subsequently lysed in 200ul nuclei lysis buffer (50mM Tris pH 8.0, 10 mM EDTA, 0.5% SDS, 1x cOmplete EDTA-free protease inhibitor) for 10 mins on a rotating wheel. The samples were then transferred to fresh 1.5ml Diagenode sonication tubes and sonicated using a bioruptor (Diagenode) by sonicating 100ul at a time (4x30’’ON-30’’OFF). After sonication, 1ml of IP dilution buffer (20mM Tris pH 8, 2mM EDTA, 0.01% SDS, 150mM NaCl, 1% Triton X-100, 1x cOmplete EDTA-free protease inhibitor) was added to each sample. The samples were then subjected to chromatin preclearing by incubating them with 25 microliters of Protein A Dynabeads (Invitrogen, Cat.N: 10002D) per sample for 2 hours on a rotating wheel. At the same time, the antibodies were conjugated with the beads by first washing 25ul of Protein A Dynabeads (Invitrogen, Cat.N: 10002D) with cold 0.5% BSA in PBS, and then adding 5ug of Neurog2 antibody (R&D Systems, Cat.N: MAB3314) and rotating for 4 hours in 500ul of 0.5% BSA in DPBS. After washing the antibody-conjugated beads with 0.5% BSA in PBS, they were added to the precleared chromatin, and incubation overnight at 4°C on a rotating wheel followed. Beads were then washed first with cold IP wash buffer I (20mM Tris pH 8, 2mM EDTA, 50mM NaCl, 1% Triton X-100, 0.1% SDS, 1x cOmplete EDTA-free protease inhibitor), two times with high salt buffer (20mM Tris pH 8, 2mM EDTA, 500mM NaCl, 1% Triton X-100, 0.1% SDS, 1x cOmplete EDTA-free protease inhibitor), one time with cold IP wash buffer II (10mM Tris pH 8, 1% NP-40, 1mM EDTA, 250mM LiCl, 1% sodium deoxycholate, 1x cOmplete EDTA-free protease inhibitor), and two times with cold TE buffer (1mM Tris pH 8, 1mM EDTA) – each wash consisted in 10 minutes of rotation. To elute DNA:Protein complexes from beads, they were incubated with 150ul of freshly made elution buffer (100mM NaHCO3, 1% SDS) for 15min at 65C (thermomixer - 800rpm), after the transfer of the supernatant, the procedure was repeated. The 300ul of DNA:protein complexes. Subsequently, 16 microliters of 5M NaCl were introduced, followed by reverse cross-linking of samples and inputs at 65°C. Afterward, they were subjected to treatment with RNase A (ThermoFisher, Cat.N: EN0531) and Proteinase K (NEB, Cat.N: P8107S), followed by purification utilizing ultrapure phenol/chloroform (ThermoFisher, Cat.N: 15593-049). Briefly, 6 μL of 10 mg/mL RNase A and the samples were incubated at 37°C for 1 hour. Subsequently, 6 μL of Proteinase K was added and the samples were incubated at 55°C for another hour. Aftwerward, an equal volume of Ultrapure Phenol:Chloroform was added, the samples were centrifuged for 10 minutes at maximum speed, the aqueous phase was then moved to a new tube and 1/10th volume of NaAc (30μL) and 1μL of glycogen were added. Subsequently, 2.5x volume of EtOH were added, the tubes were vortexed briefly and keep at -20C for 2 hours and spinned at maximum speed for 20min, washed 2x with cold 70% EtOH, and spinned again. The 70% ethanol was removed by using a pipette without disturbing the DNA pellet. The pellet was dried for 5 min at room temperature and thoroughly resuspended in 20 μL 10 mM Tris pH 7.5.

### ChIP-seq library generation

The libraries were generated by using the NEBNext® Ultra™ II DNA Library Prep Kit (NEB, Cat.N: E7103S), adhering to the manufacturer’s guidelines, with a single modification: a gel-based size selection (200-500 bp) with low met agarose (Roth, Cat.N: 6351.5) was conducted after PCR amplification. The quality of the ChIP-seq libraries was assessed using the Agilent 2100 Bioanalyser system (Agilent).

### ImmunoFACS

ImmunoFACS was performed as previously described with slight modifications (https://www.protocols.io/view/immunofacs-b2a2qage/). Dissociated NPCs were resuspended in PBS to a concentration of 1 million cells per milliliter. PFA was added to a final concentration of 1% and the samples were left to incubate for 10 minutes at room temperature with gentle rotation. To halt the reaction, a 2M glycine solution (in PBS) was added to achieve a final concentration of 0.2M, followed by a 5-minute incubation at room temperature with gentle rotation. The cells were centrifuged at 500xg for 5 minutes at 4°C, washed in cold PBS containing 1% BSA (Thermofisher, Cat.N: AM2618), re-centrifuged and then suspended in wash buffer (0.1% Saponin, 0.2% BSA Ultrapure BSA 5% (ThermoFisher, Cat. N.: AM2618), 1:100 RNAsin plus RNase inhibitor (Promega, Cat. N.: N261A), PBS) and kept rocking for 15 minutes at 4°C. PAX6-Alexa Fluor 488 (1:40; BD Biosciences, Cat.N: 561664). The sorting of G0-G1 Pax6+ cells and G0-G1 ES was performed using a FACSAria III (BD Biosciences; laser: 405 nm, 488 nm) with a 100 μm nozzle. After sorting, cells were snap-frozen and kept at -80C for subsequent 3DRAM-Seq analysis.

### Preparation of the methylation controls for 3DRAM-Seq

This procedure is performed as previously described^39^. Creating biotinylated methylation controls involved initially combining 10 μl of fully methylated pUC19 DNA (Zymo Research, Cat.N: D5017) with an equal amount of unmethylated lambda DNA (Promega, Cat.N: D1521). The mixed DNA was subjected to GpC methylation using M.CviPI methyltransferase (NEB, Cat.N: M0227), followed by purification using 1× AMPure XP beads (from Agencourt, A63881) and fragmentation to approximately 550 bp using a Covaris S220 sonicator. Next, the fragmented DNA was treated to attach biotin molecules to its sticky ends by incubating it for 6 h at 37 °C with DNA polymerase I (NEB, Cat.N: M0210) and a mixture of nucleotides containing biotin-14-dATP (Life Technologies, Cat.N: 195245016) in DpnII buffer (NEB, Cat.N: R0543S). The biotinylated DNA was then purified once again with 1× AMPure XP beads and quantified using a Qubit dsDNA HS Assay kit (ThermoFisher, Cat.N: Q32851).

### Generation of 3DRAM-Seq Libraries

The 3DRAM-Seq libraries were prepared as previously described^39^ . A detailed version of the protocol can be accessed at https://www.protocols.io/view/3dram-seq-enables-joint-epigenome-profiling-of-spa-brf8m3rw.

### Cut and Run

The Cut and Run for Rad21 in ES and NPC was conducted as previously described^14^ with 2-3.6×10^5 cells as a starting cell number.

### Cut and run library preparation

CUT&RUN libraries were prepared with the NEBNext® Ultra™ II DNA Library Prep Kit for Illumina® using 6-30 ng of fragmented DNA. The quality of the CUT&RUN libraries was assessed using the Agilent 2100 Bioanalyzer system (Agilent).

### ChIP-MS

Immunoprecipitation of Neurog2 was conducted following the “Rapid immunoprecipitation mass spectrometry of endogenous protein (RIME)” previously published protocol^62^ with some minor changes. 10×10^6 cells were collected for each replicate and fixed at room temperature for 10 minutes in ES and NPC medium containing 1% methanol-free Formaldehyde (Pierce, Cat.N: 28908) at a density of max 5 million cells/ml. Fixation was terminated by a 5-minute room temperature incubation with 0.2 M glycine. The cell pellet was washed twice with ice-cold PBS. Subsequently, cell pellets consisting of 10 million cells were promptly frozen and stored at -80°C for future use. All subsequent procedures were carried out at 4°C or on ice, with centrifugation performed at 2000 × g for 5 minutes and all the bead washes preceded by the tubes being on a magnet for 1 minute, unless otherwise specified. For each immunoprecipitation experiment, 10 million cells were utilized along with 60 μl of Protein G Dynabeads (Invitrogen, Cat.N :10004D) coupled with 3ug of the Neurog2 antibody (R&D Systems Cat.N: MAB3314).

60ul of Protein G Dynabeads (Invitrogen, Cat.N :10004D) were washed twice with LB3 (10 mM Tris-HCl pH 8.0, 100 mM NaCl, 1 mM EDTA, 0.5 mM EGTA, 0.1% Na-Deoxycholate, 0.5% N-Lauroylsarcosine) and resuspended in 100 μl of complete LB3 (LB3 containing 1 × Roche cOmplete Mini Protease Inhibitor Cocktail (Roche, Cat.N: 04693159001)). The antibody was conjugated to the beads by gentle rotation at for 1-2 hours, followed by two washes with complete LB3. Cell lysis was carried out by resuspending cells in complete LB1 (50 mM Hepes pH 7.5, 140 mM NaCl, 1 mM EDTA, 10% glycerol, 0.5% NP-40, 0.25% Triton X-100, 1 × Roche cOmplete Mini (Roche, Cat.N: 04693159001)) and incubating on a rotating wheel for 10 minutes. After centrifugation, nuclei were isolated by resuspending lysed cells in complete LB2 (10 mM Tris-HCl, pH 8.0, 200 mM NaCl, 1 mM EDTA, 0.5 mM EGTA, 1 × Roche cOmplete Mini) and incubating on a rotating wheel for 5 minutes. After centrifugation, the resulting nuclei were resuspended in 300ul of complete LB3 300 μl. Chromatin was sheared using the Bioruptor pico (Diagenode) for 6 cycles (30 seconds on/30 seconds off) at 4°C. After shearing, 1/10 volume of 10% Triton X-100 was added, followed by centrifugation at 20,000 × g for 10 minutes. The supernatant was used for chromatin immunoprecipitation and incubated with the antibody-bound beads overnight a rotating wheel. The chromatin-bound beads were washed with 4 × 1 ml of wash buffer (50 mM Hepes-KOH pH 7.6, 250 mM LiCl, 1 mM EDTA, 1% NP-40, 0.7% Na-Deoxycholate, 1 × Roche cOmplete Mini) for 10 minutes on a rotating wheel, followed by three 5-minute washes in 10mM Tris pH 8.0. Finally, the beads were transferred to a new tube and stored dry at -20°C until further analysis.

### LC-MS/MS

Beads were incubated with 10 ng/μL trypsin in 1 M urea 50 mM NH4HCO3 for 30 min, washed with 50 mM NH4HCO3, and the supernatant digested overnight (ON) in presence of 1 mM DTT. Digested peptides were alkylated and desalted prior to LC–MS analysis.

For LC–MS/MS purposes, desalted peptides were injected in an Ultimate 3000 RSLCnano system (Thermo), separated in a 25 cm analytical column (75 μm ID, 1.6 μm C18, Aurora-IonOpticks) with a 50-minute gradient from 2 to 35% acetonitrile in 0.1% formic acid. The effluent from the HPLC was directly electrosprayed into an Orbitrap Exploris 480 (Thermo) operated in data-dependent mode to automatically switch between full scan MS and MS/MS acquisition. Survey full scan MS spectra (from m/z 350 to 1,200) were acquired with resolution R = 60,000 at m/z 400 (AGC target of 3 × 106). The 20 most intense peptide ions with charge states between 2 and 5 were sequentially isolated to a target value of 1 × 105 and fragmented at 30% normalized collision energy. Typical mass spectrometric conditions were as follows: spray voltage, 1.5 kV; heated capillary temperature, 275°C; ion selection threshold, 33.000 counts.

### Library quality control and sequencing of 3DRAM-seq, Cut&Run, RNA-seq and ChIP-seq data

The libraries underwent quantificatio via qPCR employing a NEBNext Library Quant kit (New England BioLabs, Cat.N: E7630). The size distribution of these libraries was evaluated utilizing an Agilent 2100 Bioanalyzer. Subsequent sequencing was conducted either on a Illumina NextSeq1000 or a NovaSeq6000 platform.

### Bioinformatics analysis

#### Mapping of RNA-seq data

RNA sequencing libraries were sequenced with the Illumina NovaSeq 6000 platform. The reads mapped and deduplicated using STAR ^63^ with default settings. DESeq2^64^ was used to calculate fragments per kilobase of transcript per million mapped read (FPKM) and differential expressed gene values (FDR < 0.05).

#### ChIP-seq Analysis

ChIP-seq libraries were sequenced with the Illumina NovaSeq 6000 platform. The mapping of the reads onto the mm10 genome was accomplished through the chip-seq-pipeline2 from Encode with default settings. Coverage tracks in bigwig format were generated from .bam files using deepTools2^65^.

#### CUT&RUN Analysis

The Cut&Run libraries were sequenced with the Illumina Nextseq 1000 platform. The CUT&RUN data underwent standardized processing utilizing CUT&RUN tools version 2.0. Peak identification was conducted employing MACS2, while the generation of bigwig coverage tracks was accomplished utilizing deepTools2^65^.

#### 3DRAM-seq Analysis

The 3DRAM-seq libraries were sequenced with the Illumina NovaSeq 6000 platform. The mapping of the 3DRAM-seq data was performed with slightly modified version of TAURUS-MH pipeline^66^ which allowed the splitting of the reads depending on the ligation junction and the mapping of the bisulfite-converted reads with Bismark ^67^, as previously explained^39^

#### Processing of Hi-C data

The TAURUS-MH pipeline allowed the mapping of the reads and the obtaining of fragment-end-transformed read pairs, which underwent conversion in genomic ‘misha’ tracks and the subsequent import into the mm10 genonic database. The Shaman R Package ^68^ was used to shuffle observed Hi-C contacts and generate expected models, preserving overall coverage and distance distribution while removing specific features (like TADs). The Hi-C score was computed employing the k-nearest neighbors (kNN) algorithm.

#### Estimation of CpG and GpC methylation

The Bismark methylation extractor was used to determine overall CpG and GpC methylation levels by only utilizing uniquely mapped reads as input. In order to ensure a distinction between CpG and GpC methylation, the Bismark’s coverage2cytosine function contained the –nome-seq option.

#### Estimation of bisulfite conversion efficiency

We estimated the efficiency of bisulfite conversion by using Bismark in paired-end mode with the –nome-seq option (as previously described^7^) to measure the levels CpG methylation of unmethylated lambda DNA. The detection rate of methylated cytosines both in the CpG and GpC context was determined by fully methylated pUC19 DNA as well as in situ GpC methylated lambda DNA.

#### Identification of DMRs and DARs

The gNOMeHMM package was employed as previously described^39^ to identify accessible regions. Accessible peaks from the noDox and Dox conditions from the two cell types were merged to generate a unique peak dataset and methylKit was then used to determine the differentially accessible and methylated regions.

#### Contact probability analysis

Contact probability was computed as previously described^11,69,70^.

In summary, the distribution of distances between the various Hi-C contacts was analyzed by summing the observed counts per log2 bin and dividing by the total observed contacts. We measured the “contact probability scaling” exponent as the slope of the best-fit line of the cis-decay curve when plotted on log-log axes, within a chosen range of distances.

#### Insulation and TAD boundary calling

The insulation score was calculated as previously defined ^11,69,70^. Each replicate had insulation scores called separately and on the pooled contact map at 1Kb resolution within a region of ± 250Kb and was multiplied by (−1) so that high insulation score represents strong insulation. In order to account for any genome-wide changes in the insulation score, we further normalized it by multiplying with a factor defined as the average insulation score across all 1Kb genomic bins in each cell type, divided by the mean of all cell types.

We defined TAD boundaries as previously indicated^11^. Briefly, TAD boundaries were considered as the highest points within a 2kb region where the previously calculated insulation score was above the 90^th^ percentile of the analyzed genomic regions. Differential TAD boundaries were identified by using normalized insulation scores across the genome.

#### A and B compartment calling

TADs were classified as belonging to A- or B-compartments as described ^11,70^. In brief,the dominant eigenvector of the contact matrices binned at 250Kb was generated previously stated^71^, employing code available at https://github.com/dekkerlab/cworld-dekker.

The compartment strength was calculated as previously described^39^ via the log2 ratio of observed and expected inter-TAD contacts belonging to the same chromosome and at least 10Mb apart, both through the same type of domains (A-A domains, B-B domains) or intra-domain (A-B domains). It specifically represents the ratio between the sums of intra-domain contacts and the sum of inter-domain contacts.

In order to identify compartment transitions, the number of TAD boundaries between adjacent domains of different types (A-to-B domain or B-to-A domain transition) was divided by the total number of TAD boundaries within that cell type.

#### Aggregate HiC analysis

We calculated contact enrichment ratios between pairs of genomic features, such as regions identified by Neurog2 ChIP-seq peaks as previously described^7^.

In brief, we aggregated Hi-C maps to determine the log2 ratio of observed contacts compared to expected contacts within a specific window centered around the pair of genomic features being analyzed. Additionally, we measured the average enrichment of contact strength in the central part of the window and compared it to the contact strength at each of the window’s corners.

#### Average TAD and Intra- or Inter-TAD Contact Enrichment

The average TAD map was calculated as previously described^11^ .

#### TF motif analysis

We performed motif enrichment analysis using filtered list of motifs based on the JASPAR2024 database and expression in either ES or NPC (FPKM>=1) using the MonaLisa package^72^ as previously described ^14^. The k-mer enrichment analysis from figure was also generated with MonaLisa package. In order to pinpoint co-occurring TFs from the Neurog2 ChIP-seq data, we leveraged TF-COMB Python Package^73^.

#### Analysis of the proteomics data

Protein identification and quantification with iBAQ were performed using MaxQuant (version 2.4.13.0) with Database UP000000589_10090_Mmusculus_20240102; MS tol, 10 ppm; MS/MS tol, 20 ppm Da; Peptide FDR, 0.1; Protein FDR, 0.01 Min. peptide Length, 7; Variable modifications, Oxidation (M); Fixed modifications, Carbamidomethyl (C); Peptides for protein quantitation, razor and unique; Min. peptides, 1; Min. ratio count, 2. Statistical and bioinformatics analysis utilized Perseus software (version 1.5.6.0) and R. Proteins identified in the decoy reverse database or only by site modification were excluded from analysis as well as potential contaminants, and missing values were imputed based on a normal distribution.

