## Supplemental Figures for "Context-dependent epigenome rewiring during neuronal differentiation"

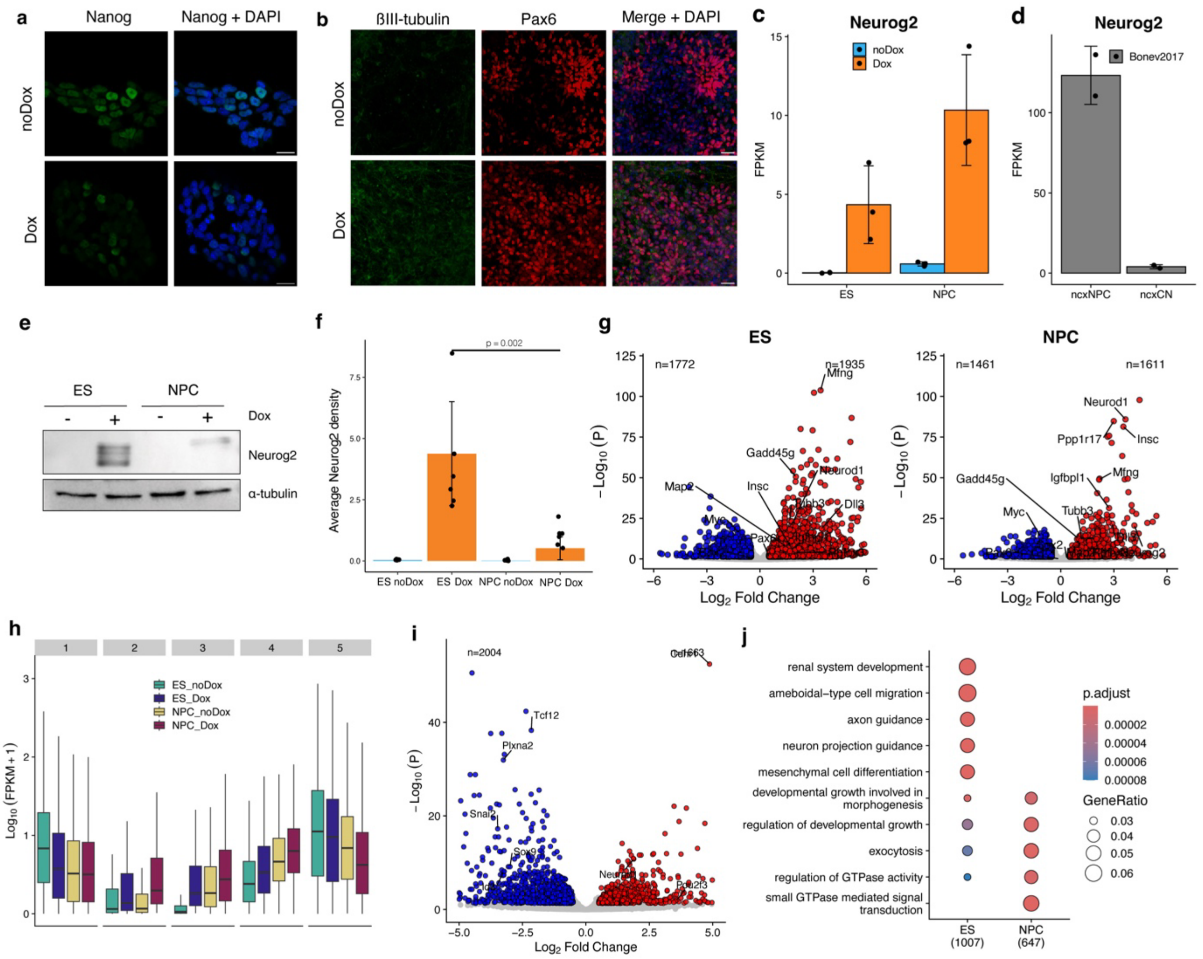


**Extended Data Fig. 1. Characterization of Neurog2 expression and induced transcriptional changes**

(a-b) Representative immunostainings of Nanog in ES cells (a), Pax6 and beta tubulin in NPC (b) with and without doxycycline supplementation. (c) Expression of Neurog2 upon Dox induction (dots represent FPKM values from individual biological replicates, error bars represent SD). (d) Expression of endogenous Neurog2 in mouse cortical progenitors (ncxNPC) or neurons (ncxCN)[11] (e) Representative western blot of whole cell lysates for Neurog2 and $\alpha$-tubulin in ES and NPCs (f) Western blot analysis and quantification of Neurog2 band intensity normalized over alpha tubulin. Values are means ± SD, n = 6 biological replicates per condition. The p value was obtained via Wilcoxon test. (g) Volcano plot displaying differentially expressed genes from a pairwise comparison between Dox vs noDox condition in ES and NPCs, respectively (n=3) (h) Boxplot displaying FPKM values for the genes in each cluster from Fig. 1F (i) Volcano plot depicting differentially expressed genes from a pairwise comparison between NPC Dox vs ES Dox conditions (n=3) (j) GO term enrichment for biological processes of the differentially expressed genes from (i).

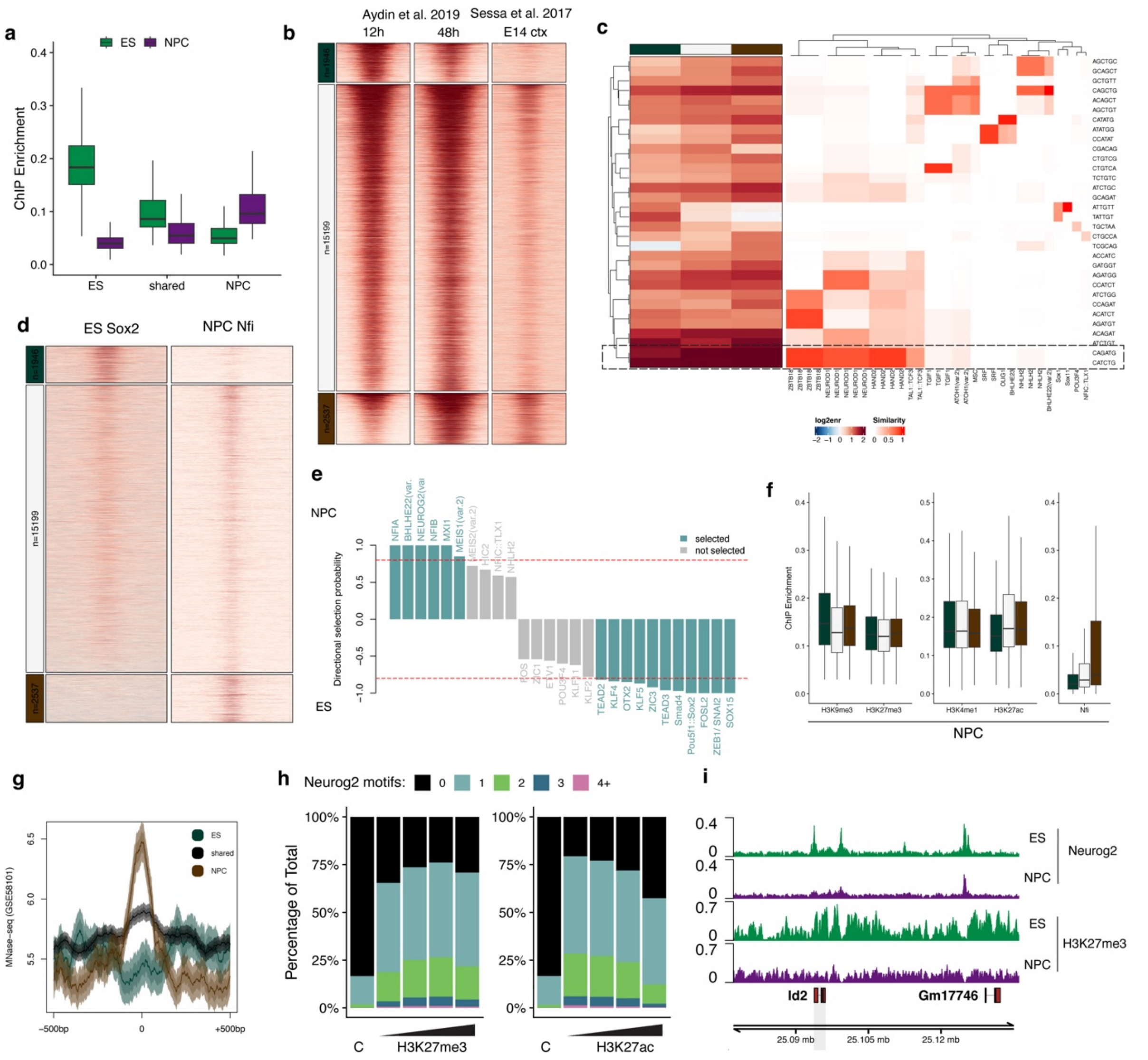

**Extended Data Fig. 2. Neurog2 cell type-specific peaks display enrichment of different sets of TFBM and pre-existing chromatin states**

(a) Boxplots depicting the ChIP-seq enrichment in reads per million (RPM) at the indicated peak groups, in ES and NPC. (b) Heatmaps showing Neurog2 ChIP-seq signal enrichment in embryoid bodies[22] and endogenous Neurog2 in mouse E14 cortex[36] (c) Heatmaps displaying kmer enrichment in the peak groups and the closest transcription factor binding motifs based on similarity. The canonical Neurog2 motif is highlighted with dashed rectangle. (d) Heatmaps showing Sox2 and Nfi ChIP-Seq peak enrichment in ES and NPC respectively centered at the same groups of peaks as Fig. 2a. (e) Directional selection probabilities for motifs identified using stability selection based on binding enrichment in NPC and ES. (f) Boxplots depicting the ChIP-seq enrichment in reads per million (RPM) of the various histone marks generated in NPCs[11] at the indicated peak categories (g) Nucleosome occupancy in ES (MNAse-seq – Carone et al., Dev Cell 2014) centered at the same groups of peaks as Fig. 2a. (h) Number of Neurog2 motifs in different peak categories, stratified by strength of H3K27me3 and H3K27ac signal in ES (i) Genomic tracks for Neurog2 ChIP-seq and H3K27me3 in both ES and NPC at the *Id2* locus.

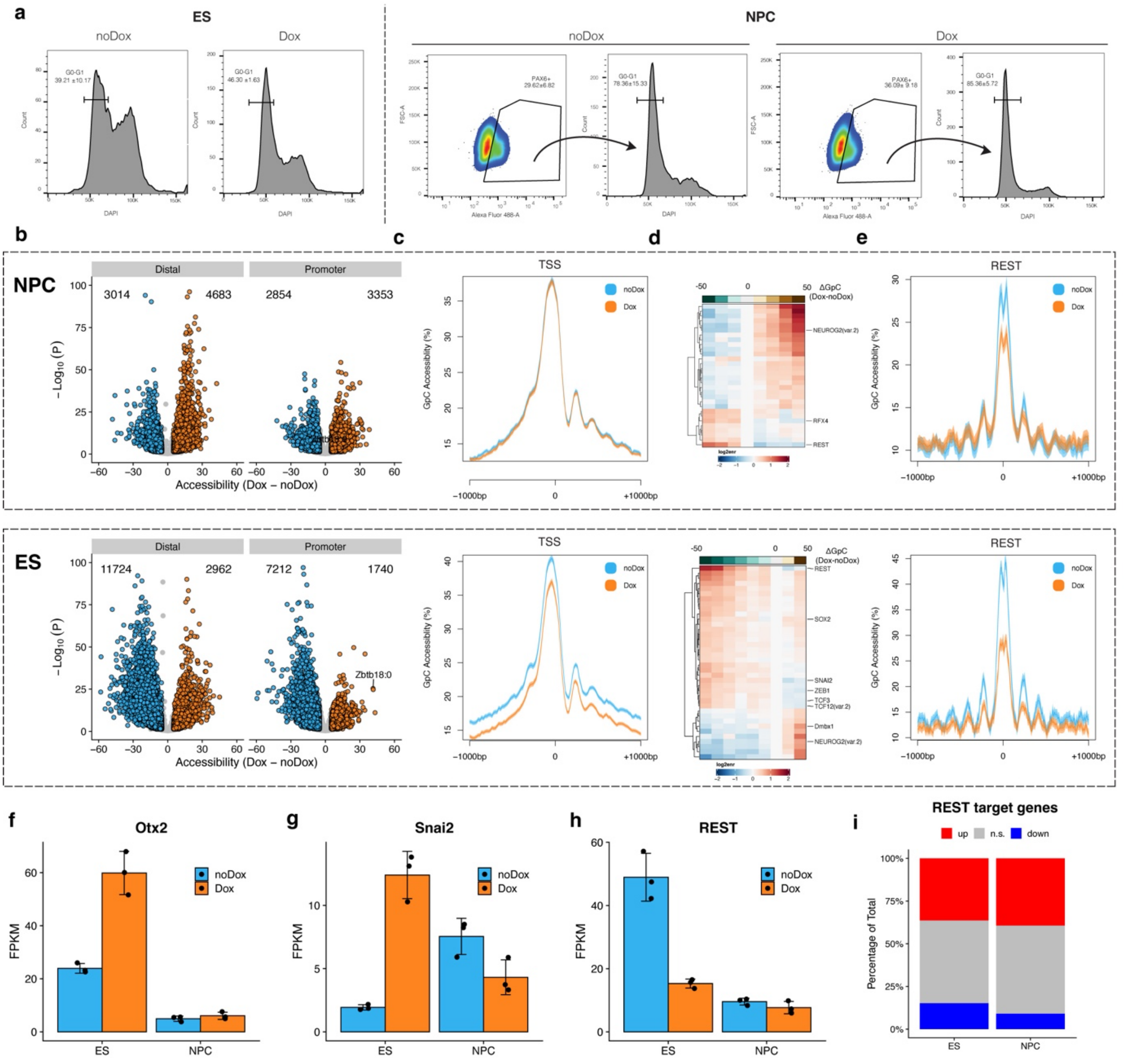

**Extended Data Fig. 3. Indirect effects on chromatin accessibility upon Neurog2 induction correlate with REST binding**

(a) Gating strategy for the immunoFACS in ES and NPC. ES cells were isolated only based on their cell cycle profile ($G_0G_1$) while NPCs were additionally selected based on the expression of the neuronal marker NPC to avoid potential confounding by differentiation heterogeneity. Numbers represent mean ± SD from the parental singlet population (b) Volcano plots displaying global differentially accessible regions (DARs) at distal and promoter regions. (c) Chromatin accessibility levels at TSS for ES and NPC. (d) Heatmap depicting motif enrichment within DARs in ES and NPC stratified by the changes in accessibility upon Neurog2 overexpression (e) Motif footprinting based on GpC accessibility levels at Rest binding sites in ES and NPCs (f-h) Expression of Otx2, Snai2 and REST in either ES or NPC (dots represent individual biological replicates, error bars represent SD). (i) Percentage of REST target genes (Johnson et al., 2008), which were upregulated, downregulated or not significantly changed (n.s.) upon Neurog2 overexpression in ES and NPC.

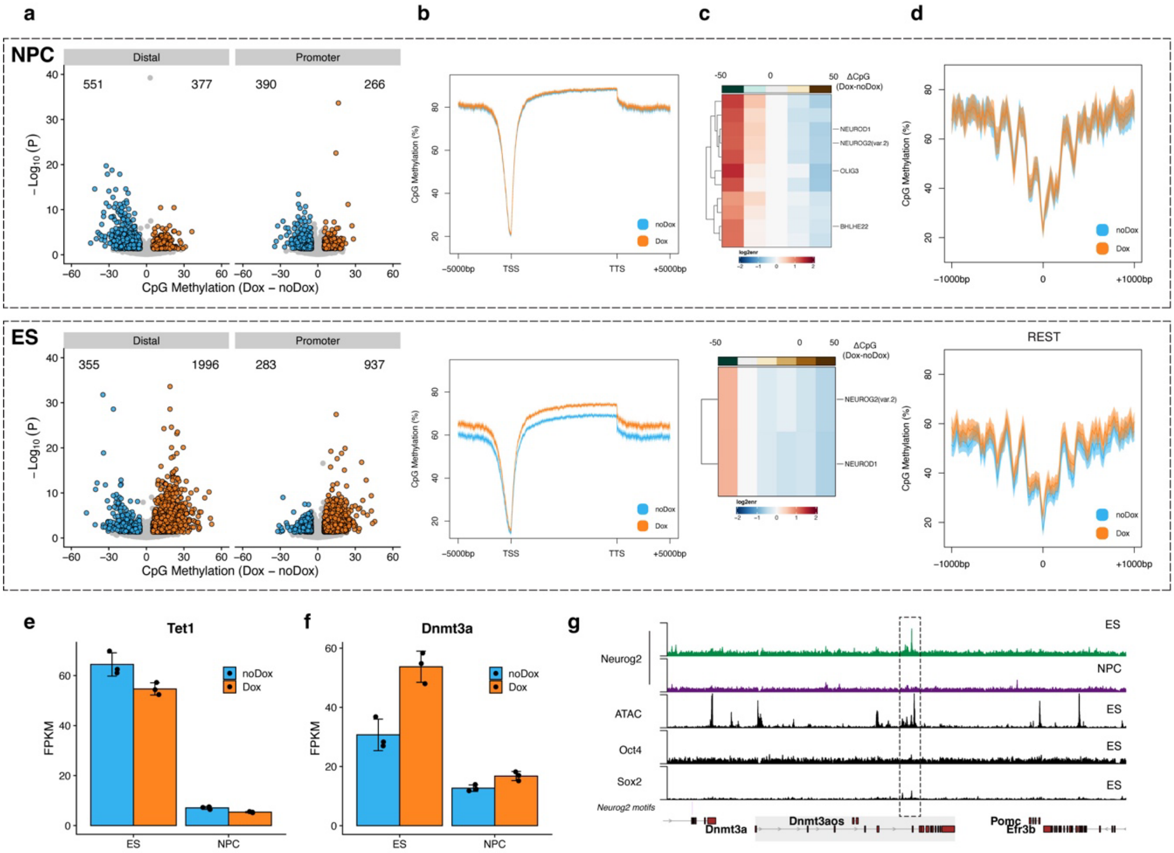


**Extended Data Fig. 4. Neurog2 leads to genome-wide increase in DNA methylation levels in ES**

(a) Volcano plots displaying global differentially methylated regions (DMR) at distal and promoter regions. (b) CpG methylation levels at gene bodies for ES and NPC (c) Heatmap depicting TFBM enrichment for DMRs in ES and NPCs stratified by the changes in methylation levels upon Neurog2 overexpression (d) DNA methylation levels at REST binding sites in ES or NPC (e-f) Expression of Tet1 and Dnmt3a in either ES or NPC (dots represent individual biological replicates, error bars represent SD). (g) Example genomic tracks depicting Neurog2, Oct4 and Sox2 ChIP-seq tracks as well as ATAC and Neurog2 motifs at the *Dnmt3a* locus. ES-specific putative enhancer co-bound by Neurog2, Oct4 and Sox2 is highlighted.

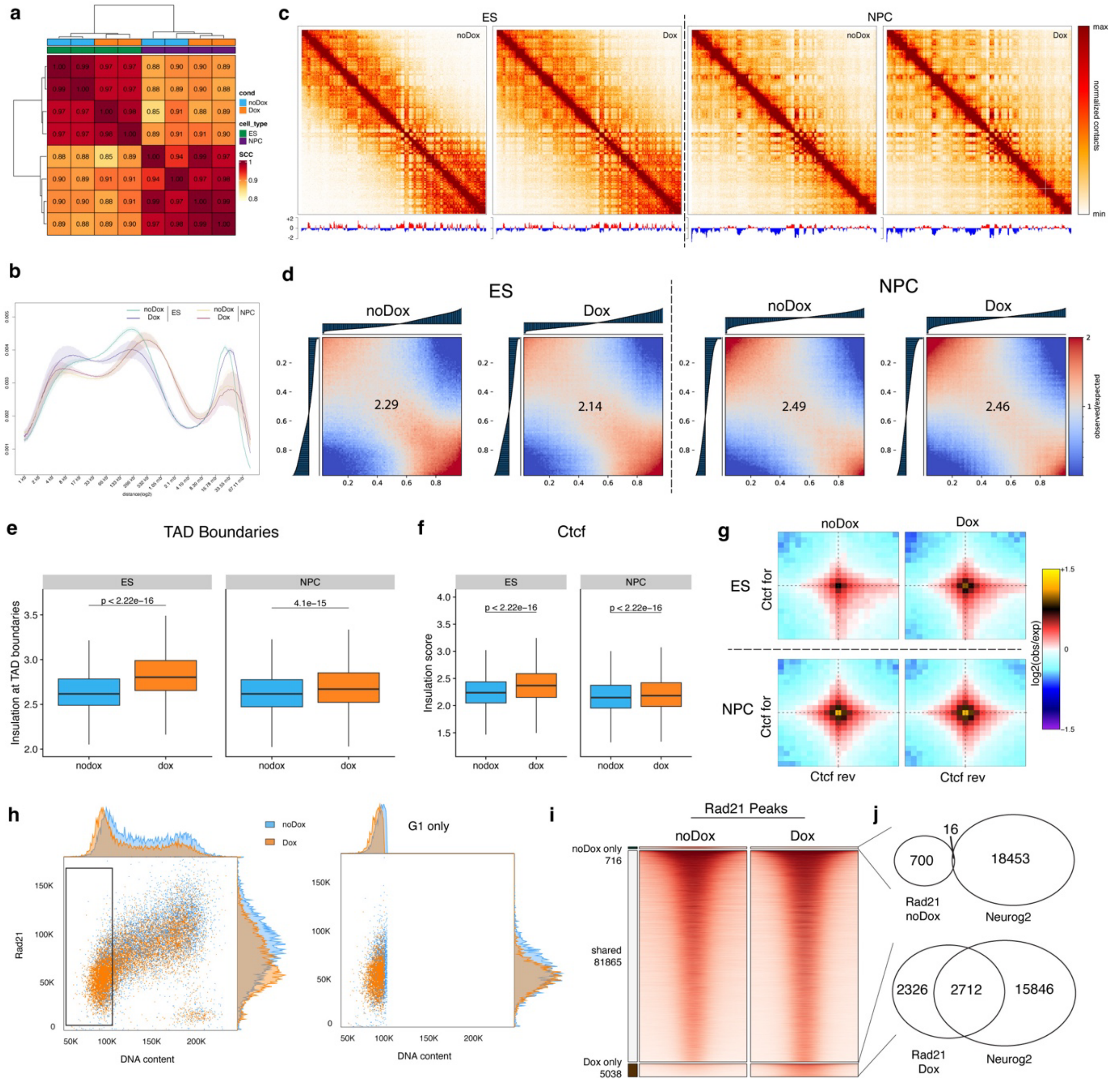


**Extended Data Fig. 5. Global 3D genome reorganization upon Neurog2 expression**

(a) Pairwise correlation matrixes displaying 3D genome correlation coefficient (stratum adjusted correlation coefficient, 25 kb bins). (b) Contact probability in logarithmic bins. Lines: mean values from biological replicates; semi-transparent ribbons: SEM (c) KR-normalized observed contact matrices for chr3 (250kb bins) and the first eigenvector (100kb bins). (d) Saddle plots showing compartment interaction strength as observed/expected contacts in 100kb bins. (e-f) Boxplots displaying insulation scores in ES and NPC at TAD boundaries (e) and (f) CTCF sites. (g) Aggregated Hi-C plots between intra-TAD pairs of convergent CTCF sites. (h) Dot plots showing the distribution of chromatin-bound Rad21 compared to DNA content using FACS. For each plot, the cell cycle profile according to DNA content appears on top while the distribution of antibody intensities is plotted on the right. (i) Heatmap displaying ES Rad21 ChIP-Seq enrichment at shared or differentially bound peaks. (j) Overlap of differentially bound Rad21 regions with ES Neurog2 peaks.
